# DeNovoCNN: A deep learning approach to *de novo* variant calling in next generation sequencing data

**DOI:** 10.1101/2021.09.20.461072

**Authors:** Gelana Khazeeva, Karolis Sablauskas, Bart van der Sanden, Wouter Steyaert, Michael Kwint, Dmitrijs Rots, Max Hinne, Marcel van Gerven, Helger Yntema, Lisenka Vissers, Christian Gilissen

**Author notes:** these authors contributed equally (Gelana Khazeeva, Karolis Sablauskas).

## Abstract

*De novo* mutations (DNMs) are an important cause of genetic disorders. The accurate identification of DNMs from sequencing data is therefore fundamental to rare disease research and diagnostics. Unfortunately, identifying reliable DNMs remains a major challenge due to sequence errors, uneven coverage, and mapping artifacts. Here, we developed a deep convolutional neural network (CNN) DNM caller (DeNovoCNN), that encodes the alignment of sequence reads for a trio as 160×164 resolution images. DeNovoCNN was trained on DNMs of 5,616 whole exome sequencing (WES) trios achieving total 96.74% recall and 96.55% precision on the test dataset. We find that DeNovoCNN has increased recall/sensitivity and precision compared to existing DNM calling approaches (GATK, DeNovoGear, DeepTrio, Samtools) based on the Genome in a Bottle reference dataset and independent WES and WGS trios. Validations of DNMs based on Sanger and PacBio HiFi sequencing confirm that DeNovoCNN outperforms existing methods. Most importantly, our results suggest that DeNovoCNN is likely robust against different exome sequencing and analyses approaches, thereby allowing the application on other datasets. DeNovoCNN is freely available as a Docker container and can be run on existing alignment (BAM/CRAM) and variant calling (VCF) files from WES and WGS without a need for variant recalling.

## INTRODUCTION

Many developmental disorders, such as intellectual disability (1), autism spectrum disorder (2) and multiple congenital anomalies (3) are known to be caused by *de novo* mutations (DNMs) (4,5). The reliable identification of DNMs is, therefore, of paramount importance both for genetic testing as well as research studies. Because of the genetic heterogeneity that exists for disorders where DNMs play a major role, the identification of DNMs is typically performed based on whole exome (WES) or whole genome sequencing (WGS) data. In principle, DNMs can be easily identified by selecting variants in the proband that are not present in either of the parents. In practice, however, this process is complicated by sequencing artifacts, mapping artifacts, differences in sequence coverage and mosaicism. Moreover, the genome of an average individual has 40 - 80 DNMs of which on average 1.45 occur in the coding regions (6), making DNMs considerably rarer than errors associated with sequencing technology. Practically this means that the sensitivity and specificity of DNM detection are usually balanced by selecting appropriate quality score cutoffs.

Several different methods have been developed to identify DNMs in next-generation sequencing (NGS) data. With methods such as DeepTrio (7) and the Genome Analysis Toolkit (GATK) (8) *de novo* calling is achieved straightforwardly by performing multi-sample variant calling and subsequent selection of variants based on genotypes corresponding to *de novo* mutations. The downside of these approaches is that DNM calling is dependent on the variant calling, which therefore always needs to be performed with the same method. For existing datasets, this may require recalling of variants with potentially high computational and storage overheads. Other tools, such as DeNovoGear (9) and TrioDeNovo (10) are able to call DNMs based on existing variant calls by modelling the probability of mutation transfer using mutation rate priors. All of these approaches provide high sensitivity, but the specificity is usually lower due to the amount of noise in NGS data, resulting in a high number of false positive calls (11). Subsequent filtering of DNMs based on quality criteria is, therefore, typically required.

Deep learning, a field of machine learning, has recently seen a growth in popularity amongst applications in genomics (12). Deep learning approaches have been able to achieve improvements in many genomics applications by converting genomic data into an image-like representation and employing convolutional neural networks (CNNs) (e.g. tumor type classification using RNA-Seq data (13) and germline variant calling (7)). Here we developed DeNovoCNN, a deep-learning model that encodes trio NGS data as images and uses a suite of CNNs to detect *de novo* mutations in next generation sequencing data.

## MATERIALS AND METHODS

### Training, validation and test datasets

A cohort of 6,067 child-parent trios was used for building the training and validation datasets, which is an extension of the cohort used in Kaplanis et. al. (5). All of the individuals were initially referred to the Radboudumc Department of Human Genetics with an indication of unexplained developmental delay, for whom trio WES was performed as described before (5). Briefly, all samples were sequenced on Illumina HiSeq 2000/4000 instruments using Agilent SureSelect v4 or v5 exome enrichment kits, respectively. Initially, *de novo* calling was performed using our in-house method based on Samtools. Subsequently, all calls from the cohort used in Kaplanis et. al. were filtered based on quality metrics as described in the original manuscript (5) and the rest of the cohort was filtered according to the following approach: GATK quality score > 300 for substitutions and > 500 for insertions and deletions, coverage ≥ 20X in the proband, VAF > 30%. The complete dataset yielded 13,068 DNM calls, which were used to construct the training and validation datasets (Supplementary Figure 1).

Snapshots of all of the potential DNM calls were generated using the Integrative Genomics Viewer (IGV) (14) for visual inspection, and each variant was evaluated by assigning it to one of the three classes: DNM, IV (inherited variant) or UN (unknown) for cases where it was not feasible to make the confident decision on visual inspection alone. UN variants were removed from the dataset. The obtained dataset of 5,616 trios was complemented with randomly selected IVs resulting in 10,274 DNMs and 55,134 IVs. The 5,616 trios were randomly divided into training, validation and test subsets using a 70/15/15 percentage ratio. (Supplementary Figure 1).

One of the challenges of DNM detection is to distinguish false positives in difficult genomic regions, so we developed a way to add such examples. First, we took the current training and validation datasets to train an interim DeNovoCNN model for DNMs calling. Second, we randomly selected 403 trios and applied an interim DeNovoCNN model to get candidate DNM calls on these trios. Finally, we manually curated all calls in IGV, selecting 905 IVs that were either clearly inherited from the parents or occurred in difficult regions where a lot of sequencing mistakes and artefacts were visible. In addition, we selected 159 true DNMs. This provided a better representation of the locations where our algorithm made mistakes in the previous step and therefore the most difficult genomic regions for the model.

Despite the large exome dataset, the total number of DNMs for training was relatively low. Therefore, we supplemented the DNM dataset by performing DNM calling using the in-house caller on 2 artificial trios where the child was unrelated to the parents. These 2 trios were constructed by randomly sampling 2 parent pairs, followed by the random choice of a child (Supplementary Figure 1). This resulted in an additional 1,005 DNMs that were added only to the training dataset to avoid biases in validation and test datasets.

All IVs and DNMs were further assigned into three categories: insertions, deletions and single-nucleotide substitutions for the training of the three different models. (Supplementary Table 1).

### DeNovoCNN

#### Model architecture

We aimed to replicate the visual inspection process of possible DNMs performed by human experts using software such as IGV. By converting NGS data into RGB images *de novo* variant calling could be approached as a computer vision classification task with two classes: DNMs and IVs. The state-of-the-art approach for vision classification tasks is the convolutional neural network (CNN), a variation of which we chose for our purposes. The choice of the architecture was a trade-off between the ability to generalize (complexity) of the model and the available amount of training data. Thus, the model architecture was chosen to be basic and consisted of 9 2D convolutional layers with 96 filters, 3x3 kernels, ReLU activation and the same padding in each layer. After every third convolutional layer, we applied batch normalization and added a Squeeze-and-Excitation block (15). Global max and average pooling were applied before the output layer (Supplementary Figure 2). The architecture was developed using Python with the TensorFlow v.2.3.0 (16). Using this architecture we constructed three separate models, for insertions, deletions and substitutions because of their specific visual patterns and the skewness of the dataset towards substitutions. We also considered a single model for all three types of variants but obtained inferior results using this approach.

#### Image generation

Variants in *de novo* and control datasets were converted into images prior to being fed to the convolutional neural network. All variants of interest were converted into 160×164 RGB images (Supplementary Figure 3). Image generation was based on reads pileup data in the location of the variant capturing 20 nucleotides before and after the candidate DNM. Read pileup data from individual trio members for the same variant position was extracted using the Pysam v.0.19.0 library (17).

Each row in the image encodes a read base sequence. Image columns were structured in a recurring pattern of 4 pixels per genomic position, which represents a one-hot vector that encodes A, C, T and G bases respectively. Thus, the image width of 164 pixels represents a sequence of 41 (164 / 4) bases with the variant starting at the central position (20 using 0 indexing). In a one-hot vector for (A, C, T, G) the coordinate was filled with a value in the resulting image in case we observe this nucleotide in the corresponding genomic position in the read, whereas the rest were filled with zeros (Figure 1). Pixel intensities have a maximum value of 255, adjusted by mapping and base quality scores with higher quality corresponding to higher pixel intensity. Each column represents the sequencing depth which was limited to 160 reads for computational performance. Red, green, and blue color channels represent different individuals of the trio, corresponding to child, father, and mother respectively.

**Figure 1.**
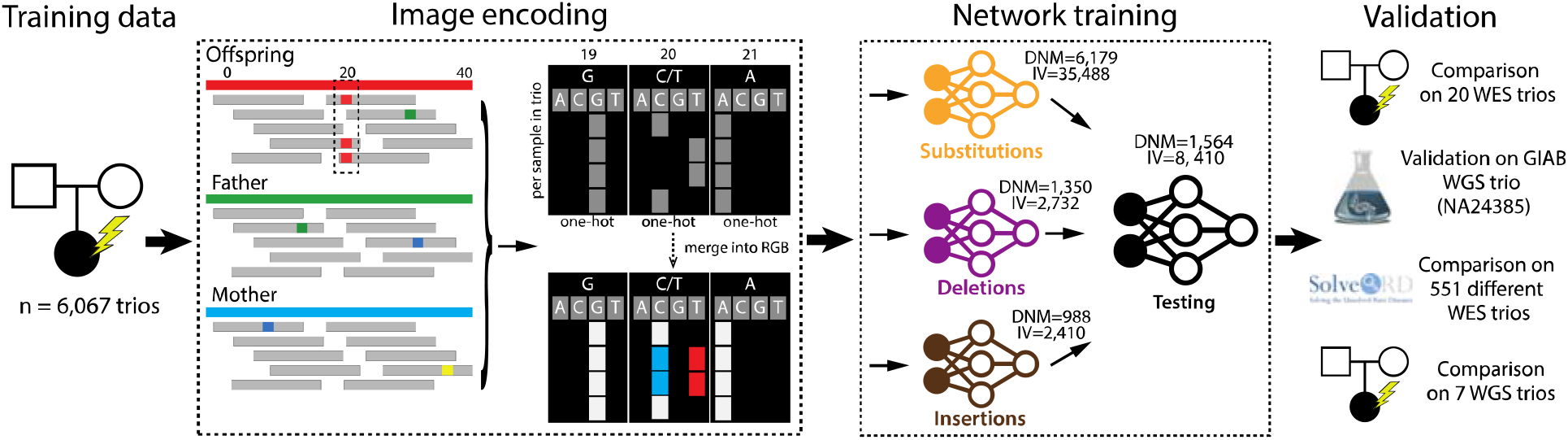
Overview of the method depicting the different steps starting with (from left to right), the training data, encoding of sequence data as images, training of the different deep-learning models, and validation of the final model. The first step consisted of the construction of training and validation datasets using *de novo* and inherited or false *de novo* variants from the 6,067 trios. As is shown in the “Image encoding” section of the figure all variants were transformed to RGB images, where each color channel corresponds to the child, father, and mother data respectively. Every row encodes a separate read, and every nucleotide in a particular position is encoded as a one-hot vector, containing (A, C, T, G). 70% of these images were then used to train 3 convolutional neural networks (for substitutions, deletions, and insertions variants). The remaining 15% of the data (1,564 DNM variants, 8,410 IV variants) was used as a test dataset. The resulting models were thoroughly validated using various independent datasets: the external gold standard GIAB WGS trio, the in-house 20 WES dataset, the in-house 7 WGS dataset, and the multi-platform 551 WES trios from the SolveRD project.

#### Hyperparameters optimization

The architecture of the model and the process of training require the definition of some hyperparameters, such as learning rate, number of convolutional features, batch size, and regularization coefficients. The choice of these parameters was done using the Hyperband algorithm for hyperparameter optimization (18) (Supplementary Table 2). The values for the number of convolutional features and batch size were sampled from [32, 64, 96, 128] and [32, 64] respectively. For continuous parameters the values were logarithmically sampled from corresponding segments. The L1 coefficient of the sigmoid layer was sampled from [1e-10, 0.1], learning rate from [1e-8, 0.01] and the Adam weight decay from [1e-8, 0.01]. The Hyperband optimization was performed such that the hyperparameters showed the lowest cross-entropy loss on the validation dataset.

#### Training the model

Networks were trained for 100 epochs unless the performance on the validation dataset did not improve for 40 epochs, in which case the training was stopped. For all 3 networks training stopped before reaching 100 epochs (Supplementary Figure 4). The final models were selected at the epoch that showed the best performance on the validation dataset. Due to the large dataset size, the substitution network was trained first using random weight initialization, while insertion and deletion networks were trained using weights from the trained substitution network as the starting point. As a result of optimization, some hyperparameters are different for the three different networks (Supplementary Table 2). Adam optimizer for substitutions and AdamW for insertions and deletions with default Keras parameters were used for minimization of binary cross-entropy loss in all models. The initial learning rate was set to the optimized values for each network with a stepwise decay of 0.5 every 10 epochs (Supplementary Table 2). The output of the network is a vector containing probabilities for a variant being a DNM and IV. The area under the curve (AUC), overall accuracy, specificity, sensitivity and F1 score were calculated on the test set.

Data augmentation was applied during the training of the networks for substitutions, deletions and insertions. The standard augmentations included random brightness adjustment by a factor between [0.3, 1]. To increase the stability of the model, random shuffling of the reads was implemented. Additionally, we observed that some specific cases of DNMs were underrepresented in our dataset, namely, DNMs in low-coverage regions and multi-nucleotide substitutions. We simulated reduced coverage by discarding a random number of reads from the pileup and enriched the dataset for multi-nucleotide substitutions by generating adjacent substitutions using the substitution dataset on-the-fly.

The training was performed on a machine with NVIDIA GeForce GTX TITAN X 12 Gb. Training time on this machine was approximately 17.6 hours for substitutions, 1.7 hours for deletions, and 1.5 hours for insertions.

#### DNM prediction

After the training DeNovoCNN was used for DNM prediction on new data. The input consists of one VCF and one BAM file per sample (three per trio), as well as paths to the trained model weights for substitutions, insertions and deletions. The prediction consists of two steps. The first step is dependent on the initial variant calling used for generating VCF files: using the trio VCF files, the inherited variants are discarded. This is achieved using the `bcftools isec -C child_vcf father_vcf mother_vcf` (17) command and results in around a 10-fold reduction of the number of genomic locations for evaluation, which usually ends up with <10,000 variants for WES (depending on capture kit size) and ∼100,000 for WGS. The second step iterates through the generated list and classifies each variant as DNM or non-DNM. The variant is considered to be *de novo* if the probability of DNM class returned by DeNovoCNN is higher or equal to 0.5. The application of DeNovoCNN doesn’t require any GPUs. On a standard 16 core CPU machine, the run-time is approximately 15 – 20 minutes for a 100x coverage WES trio and 5.5 - 6 hours for a 50x WGS trio.

#### Performance assessment

In order to validate the performance of the proposed deep learning model, we used datasets from different types of sequencing (exomes and genomes) as well as different types of enrichment and sequencing platforms (Supplementary Table 3, Supplementary Methods). DeNovoCNN was compared to other available algorithms, such as DeepTrio (7), GATK PossibleDeNovo (8), and DeNovoGear (9). We also compared to our in-house *de novo* detection algorithm, based on Samtools mpileups, because of the 10 years of experience that we had with this approach.

For validation purposes, we applied the above-mentioned algorithms as follows:

- DeNovoCNN takes as an input trio VCF files and BAM/CRAM files. See the DNM prediction section for the details.
- DeepTrio version 1.2.0 was run on BAM/CRAM files to call variants on a trio, followed by GLnexus tool for gVCF merging according to DeepVariant and GLnexus best practices (19) with optimized configuration for DeepVariant caller in WES and WGS data (DeepVariantWES, DeepVariantWGS). We additionally ran GLnexus using a config with no quality filters (DeepVariant_unfiltered) as it is suggested in the DeepTrio available documentation. Since it is recommended not to perform BQSR on the input files for DeepTrio, we run the tool on the BAMs without base recalibration as well. *De novo* mutations were defined as Mendelian violations with a heterozygous variant in the child and homozygous reference calls in the parents and were selected with the RTG Tools Mendelian package v.3.12.1 (20). Whereas DeepTrio performed well on WGS data, we were unable to generate good results for DeepTrio on WES data. Although we tried different settings and post-analysis filtering, we found that DeepTrio consistently generated high numbers of DNM calls in WES data. We have documented our efforts and results in the Supplemental (Supplementary Table 4).
- DeNovoGear version 1.1.1 (9) was run according to the specifications. DeNovoGear takes in PED and BCF files as input. The BCF file was generated using the following command: `samtools mpileup -gDf reference.fa child.bam father.bam mother.bam`
- GATK (gatk4-4.1.2.0 and gatk4-4.1.8.1) was run on BAM/CRAM or gVCF files according to the best practices for germline short variant discovery (SNPs + indels) and “Genotype Refinement workflow for germline short variants” (8) to detect *de novo* variants.
- Our in-house method: this is an in-house developed method that generates a list of *de novo* candidates based on the VCF files of the trio (1). For WES remaining candidates were then filtered out based on gnomAD allele frequency <1.0%. For WGS data gnomAD allele frequency <0.1% and xAtlas quality score >15 filters were used. Subsequently, this method performs Samtools pileups which are used to select the most likely DNM candidates based on a set of hard filters. The variant is considered to be *de novo* if the coverage in both parents is ≥10 and either there are no alternative reads in the parents or the VAF in both parents ≤15% with the number of alternative reads <3.

We compared these methods based on the raw *de novo* calls, as well as after the application of the commonly used quality filtering for *de novo* mutations. This high-quality DNM set was created using the following filters (derived from in-house practices): number of reads in the sample and both parents ≥10, variant allele frequency ≥20%, gnomAD allele frequency <0.01%.

## RESULTS

We used a convolutional neural network (CNN) with squeeze-and-excitation blocks (15) as the architecture for DeNovoCNN (Figure 1, Supplementary Figure 2), and trained three separate models for substitutions, insertion and deletion variants. Our primary training dataset was based on DNMs and inherited variants (IVs) from a cohort of 6,067 child-parent trios that were supplemented with simulated data in order to optimize training (5), (Supplementary Figure 1, Material and Methods). All variants were converted into 160×164 RGB images (Supplementary Figure 3). The dataset was split into an 70% training (8,517 DNMs, 40,590 IVs), 15% validation (1,357 DNMs, 7,110 IVs; Supplementary Table 1) and 15% test (1,564 DNMs, 8,410 IVs) dataset. DeNovoCNN generates a probability of a variant being *de novo*, and, therefore, we used a threshold of ≥0.5 to select *de novo* calls. After training DeNovoCNN achieved a high sensitivity/recall rate of 96.74% (substitutions: 97.71%, insertions: 91.76%, deletions: 91.76%) and precision of 96.55% (substitutions: 97.78%, insertions: 96.3%, deletions: 87.64%) (Supplementary Figure 5, Supplementary Figure 6, Supplementary Table 5) on the test dataset.

### Comparison on GIAB WGS dataset

In order to compare our method to other *de novo* detection methods on an independent dataset, we used Illumina WGS data of an Ashkenazim Trio (NA24385; NA24149; NA24143) from the Genome in a Bottle (GIAB) consortium (21). This trio was sequenced using various different technologies in order to create a dataset of 1,323 high-quality cell-line and germline DNMs. We only considered DNMs in high-quality regions, as suggested by the GIAB consortium. We compared DeNovoCNN DNMs to DNMs from DeepTrio (applied on BAMs with and without BQSR, with DeepVariantWGS and DeepVariant_unfiltered presets for GLnexus), in-house method based on Samtools (17), GATK (8), GATK filtered for high confidence DNM calls (GATK_HC), and DeNovoGear filtered using a ≥0.5 and ≥0.9 probability thresholds (DeNovoGear-0.5, DeNovoGear-0.9) (9) (Figure 2A, Table 1, Supplementary Table 6, Supplementary Figure 7).

**Figure 2.**
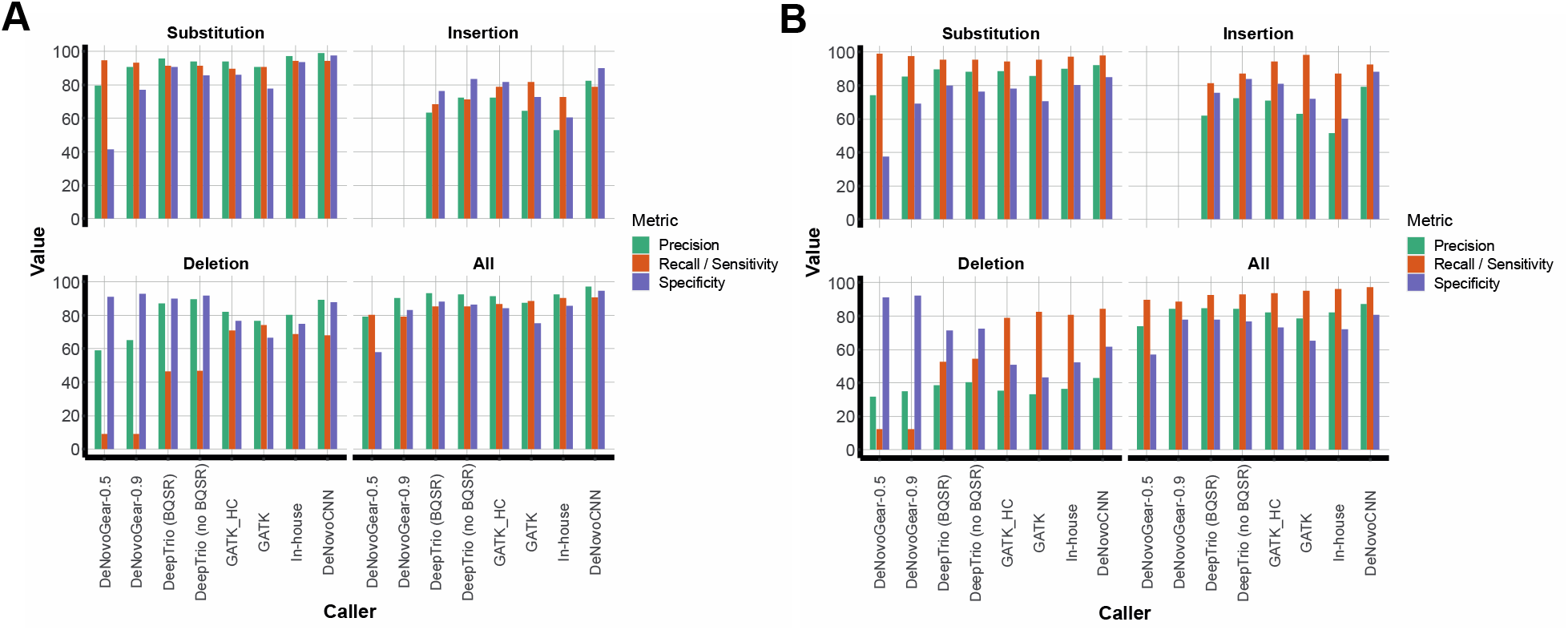
The comparison results on the GIAB reference trio. These figures contain 4 plots of the different tools’ performance for substitutions, insertions, deletions, and all variants on a GIAB dataset. All tools for the comparison are on the horizontal axis: DeNovoGear with a threshold of 0.5, DeNovoGear with a threshold of 0.9, DeepTrio (DeepVariant_unfiltered, BQSR), DeepTrio (DeepVariant_unfiltered, no BQSR), GATK with high quality calls only, GATK, our in-house tool, and DeNovoCNN. The green, orange and violet bars show precision, recall (sensitivity), and specificity respectively. A. The performance of different *de novo* calling methods for the GIAB reference genome trio. B. The performance of different *de novo* calling methods for the GIAB reference genome trio using the manually curated set of DNMs (Material and Methods).

**Table 1.** The results of the comparison on the GIAB dataset. Every row shows different statistics and performance metrics for DeNovoCNN, DeNovoGear with the probability threshold of 0.5 and 0.9, DeepTrio with different settings, GATK and GATK with high quality DNMs, and our in-house tool. The first column shows the name of the tool, the next five columns show the number of total DNM calls, the number of true positive, false positive, true negative and false negative DNM calls respectively based on GIAB reference calls. The next 4 columns show the performance metrics, such as sensitivity/recall, specificity, precision and accuracy.

| Tool | Calls | TP | FP | TN | FN | Sensitivity/<br>Recall | Specificity | Precision | Accuracy |
| --- | --- | --- | --- | --- | --- | --- | --- | --- | --- |
| DeNovoCNN | 1233 | 1198 | 35 | 640 | 125 | 90.55 | 94.81 | 97.16 | 91.99 |
| DeNovoGear-0.5 | 1346 | 1063 | 283 | 392 | 260 | 80.35 | 58.07 | 78.97 | 72.82 |
| DeNovoGear-0.9 | 1161 | 1047 | 114 | 561 | 276 | 79.14 | 83.11 | 90.18 | 80.48 |
| DeepTrio (DeepVariantWGS, BQSR) | 210 | 176 | 34 | 641 | 1147 | 13.3 | 94.96 | 83.81 | 40.89 |
| DeepTrio (DeepVariantWGS, no BQSR) | 268 | 240 | 28 | 647 | 1083 | 18.14 | 95.85 | 89.55 | 44.39 |
| DeepTrio (DeepVariant_unfiltered, BQSR) | 1207 | 1127 | 80 | 595 | 196 | 85.19 | 88.15 | 93.37 | 86.19 |
| DeepTrio (DeepVariant_unfiltered, no BQSR) | 1222 | 1129 | 93 | 582 | 194 | 85.34 | 86.22 | 92.39 | 85.64 |
| GATK | 1338 | 1171 | 167 | 508 | 152 | 88.51 | 75.26 | 87.52 | 84.03 |
| GATK_HC | 1257 | 1149 | 108 | 567 | 174 | 86.85 | 84.0 | 91.41 | 85.89 |
| In-house | 1293 | 1195 | 98 | 577 | 128 | 90.33 | 85.48 | 92.42 | 88.69 |

DeNovoCNN outperformed other algorithms with a precision rate of 97.16%. DeepTrio (DeepVariant_unfiltered, BQSR) showed a precision of 93.37%, our in-house tool showed a precision of 92.42%, and DeepTrio (DeepVariant_unfiltered, no BQSR) showed 92.39%, while GATK_HC and DeNovoGear-0.9 performances were 91.41% and 90.18% respectively. DeNovoCNN also has the highest sensitivity/recall rate of 90.55%, our in-house tool showed 90.33%, and DeepTrio (DeepVariant_unfiltered, no BQSR) showed 85.34%, GATK and DeNovoGear-0.5 performance were 88.51% and 80.35% respectively (Figure 2A, Table 1). We note that DeepTrio was actually trained on this GIAB dataset, and therefore the DeepTrio results may be slightly inflated.

We observed a relatively low recall for insertions and deletions of all algorithms which led us to investigate possible problems with the GIAB high-quality DNM dataset (Supplementary Table 6). Therefore, we performed manual cleaning of likely false positive *de novo* insertion and deletion calls in the GIAB gold standard dataset based on visual inspection (Supplementary Methods). We then repeated our comparison on this manually curated GIAB dataset and found that DeNovoCNN still outperformed all other methods on sensitivity/recall as well as precision and accuracy (Figure 2B, Supplementary Table 7).

### Comparison on 20 WES trios

The comparison on the GIAB dataset highlighted the difficulties in obtaining high-quality validation datasets for DNMs. Therefore, we also compared DeNovoCNN performance with GATK_HC, DeNovoGear, and our in-house tool on a dataset of 20 randomly selected in-house WES trios that were not part of the original training dataset of 6,067 trios (Figure 3A, Table 2, Material and Methods). This allowed us to validate the called DNMs experimentally. For all of the DNM calls by the different methods we performed a visual selection to discard obvious false-positive variants. The remaining 50 variants were validated by Sanger/IonTorrent sequencing of which 24 were confirmed as DNMs, and 4 were either inherited or false positives (Supplementary Figure 8). For 22 variants it was not possible to perform the validation due to difficulties with designing the primers (9 variants) and depletion of the DNA sample (13 variants). Based on these validations, DeNovoCNN was able to correctly identify all 24 confirmed DNMs (DeNovoCNN sensitivity/recall is 100%, next best is 83.33% for in-house, GATK and GATK with high quality calls, and DeNovoGear-0.9 has 79.17%), showing the best performance on all other metrics as well (Table 2). We subsequently performed additional quality filtering on the DNMs (Materials and methods), as is likely to happen in a real-world setting, and re-evaluated the results. DeNovoCNN still showed the best results on all calculated metrics (Supplementary Table 8).

**Figure 3.**
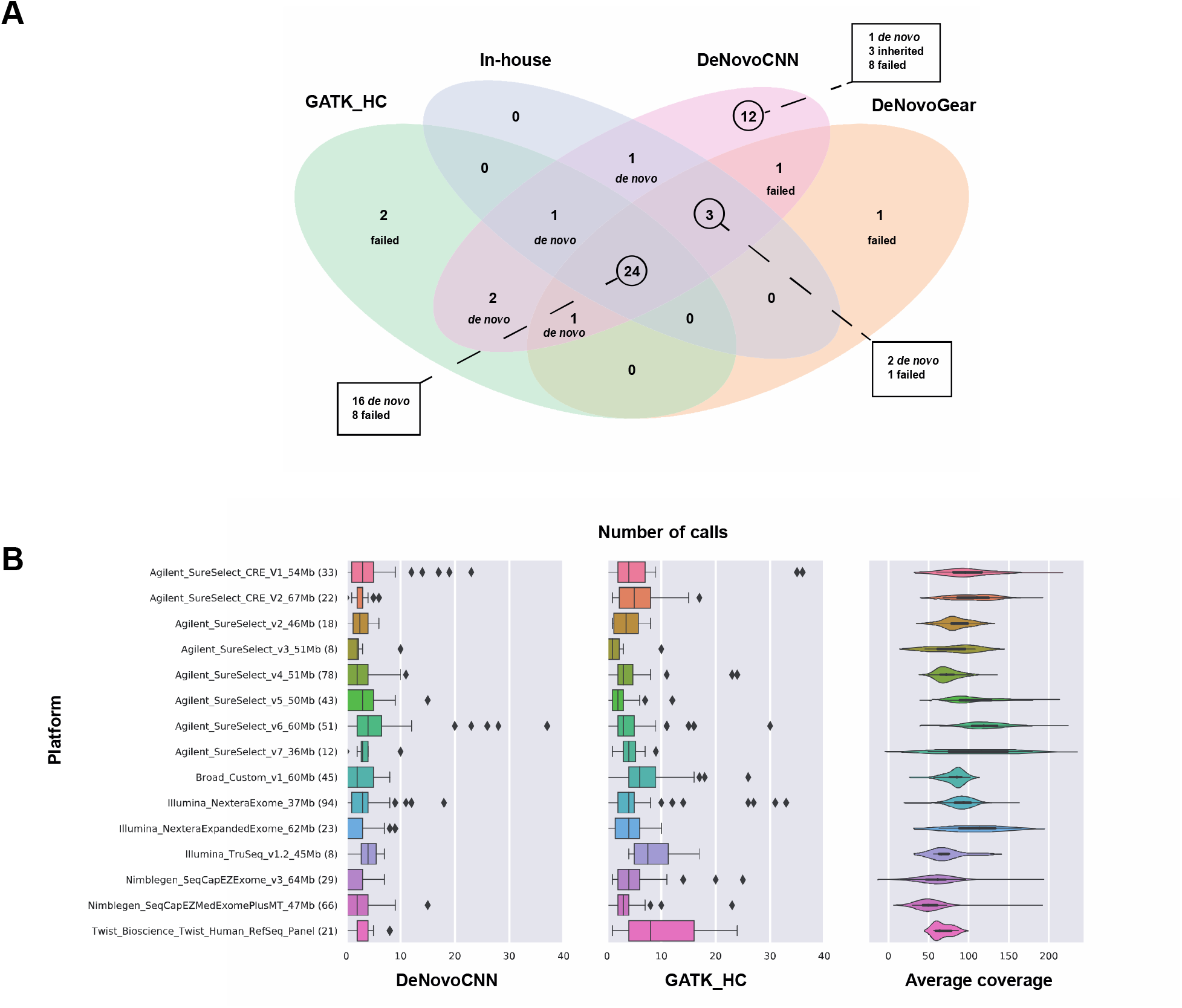
Validation and comparison of the DeNovoCNN method with other tools on exome data. **A. Comparison results on 20 in-house WES trios based on DNMs that were validated by Sanger/ IonTorrent sequencing**. The graph contains only variants that were marked as potentially *de novo* after manual validation in IGV and then sent for Sanger/IonTorrent validation (see Materials and Methods). The green oval contains all potentially *de novo* variants for the GATK tool with high quality DNMs, the light blue oval indicates potential *de novo* variants according to the in-house tool, the magenta oval contains potentially *de novo* variants according to the DeNovoCNN tool, and finally, light brown oval shows potentially *de novo* variants based on the DeNovoGear tool. Each intersection of the circles contains the number of potentially *de novo* variants that were found in the overlap of the corresponding tools’ calls. For each intersection, the results of the Sanger validation of these variants are shown. **B. The DeNovoCNN and GATK results on SolveRD samples from different exome sequencing kits and platforms**. For each sequencing kit, the number of samples used is indicated in brackets. The first two graphs show the distribution of the number of calls (on the horizontal axis) per sequencing kit using boxplots for DeNovoCNN and GATK with high-quality DNMs respectively. The last graph shows the distribution of the average coverage per sequencing kit using violin plots.

**Table 2.** The results of the Sanger/IonTorrent validations on the 20 WES trios based on raw calls of the tools. Every row shows different statistics and performance metrics for DeNovoCNN, GATK with high quality DNMs and GATK, DeNovoGear with a probability threshold of 0.9, and our in-house tool. The first column shows the name of the tool, the next five columns show the number of total DNM calls, the number of true positive, false positive, true negative and false negative DNM calls respectively based on the results of Sanger/IonTorrent validations. The next 4 columns show the performance metrics, such as sensitivity/recall, specificity, precision and accuracy.

| Tool | Calls | TP | FP | TN | FN | Sensitivity/<br>Recall | Specificity | Precision | Accuracy |
| --- | --- | --- | --- | --- | --- | --- | --- | --- | --- |
| DeNovoCNN | 75 | 24 | 51 | 2488 | 0 | 100.0 | 97.99 | 32.0 | 98.01 |
| GATK_HC | 103 | 20 | 83 | 2456 | 4 | 83.33 | 96.73 | 19.42 | 96.61 |
| GATK | 147 | 20 | 127 | 2412 | 4 | 83.33 | 95.0 | 13.61 | 94.89 |
| DeNovoGear-0.9 | 599 | 19 | 580 | 1959 | 5 | 79.17 | 77.16 | 3.17 | 77.18 |
| In-house | 85 | 20 | 65 | 2474 | 4 | 83.33 | 97.44 | 23.53 | 97.31 |

Additionally, we compared different properties (VAF, coverage, strand bias, variant context and population allele frequency) between the true positive and false positive DNMs from DeNovoCNN, GATK and DeNovoGear in order to explore obvious possibilities for improvements (Supplementary File). We did not observe any striking differences, with the exception of population allele frequency (gnomAD) which could be used as post-processing filtering.

### Results on the multi-platform WES trio dataset

Next, we wanted to verify that DeNovoCNN is robust across different capture and sequencing approaches. We used an exome dataset of the Solve-RD consortium (22) that contains 551 trios sequenced across 15 different capture/sequencing combinations (Supplementary Methods; Supplementary Table 9). We measured the robustness of our method by considering the number of called DNMs per sample and compared this to the number of high-quality DNM calls from GATK (GATK_HC) (Figure 3B, Supplementary Figure 9). In addition, we expected that the number of calls is within the same range regardless of the sequencing platform that was used. The median number of DeNovoCNN calls is 2.5, the 5th and 95th percentiles are 0.0 and 7.55 whereas the overall distribution lies between 0 and 64 calls. This result is consistent with what we observe for GATK calling (median number of calls is 4.0, the 5th and 95th percentiles are 1.0 and 13.0, maximum number of calls is 93.0). This suggests that our method was likely not overfitted to the training dataset’s specific capture kit and sequencing instrument.

### Results on 7 WGS trios

To confirm that DeNovoCNN also performs well on whole-genome sequencing data, we used 7 in-house WGS trios. We applied DeNovoCNN, GATK and DeepTrio (with DeepVariantWGS and DeepVariant_unfiltered presets for GLnexus). We compared these tools with high-quality *de novo* calls obtained with PacBio Hi-Fi long reads sequencing (LRS) (Supplementary Methods). DeNovoCNN had the highest concordance with LRS calls with sensitivity/recall of 81.83%, next was GATK with 76.37% and 75.72% and for low and high confidence calls respectively. DeepTrio had 73.47% and 15.76% concordance with DeepVariant_unfiltered and DeepVariantWGS preset filtration respectively. DeNovoCNN outperformed other tools significantly based on the specificity (92.81%), precision (21.8%), and accuracy (92.54%) (Figure 4, Table 3). We repeated the comparison after filtering the DNMs for high quality calls (Materials and methods) and obtained similar results (Figure 4, Supplementary Table 10).

**Figure 4.**
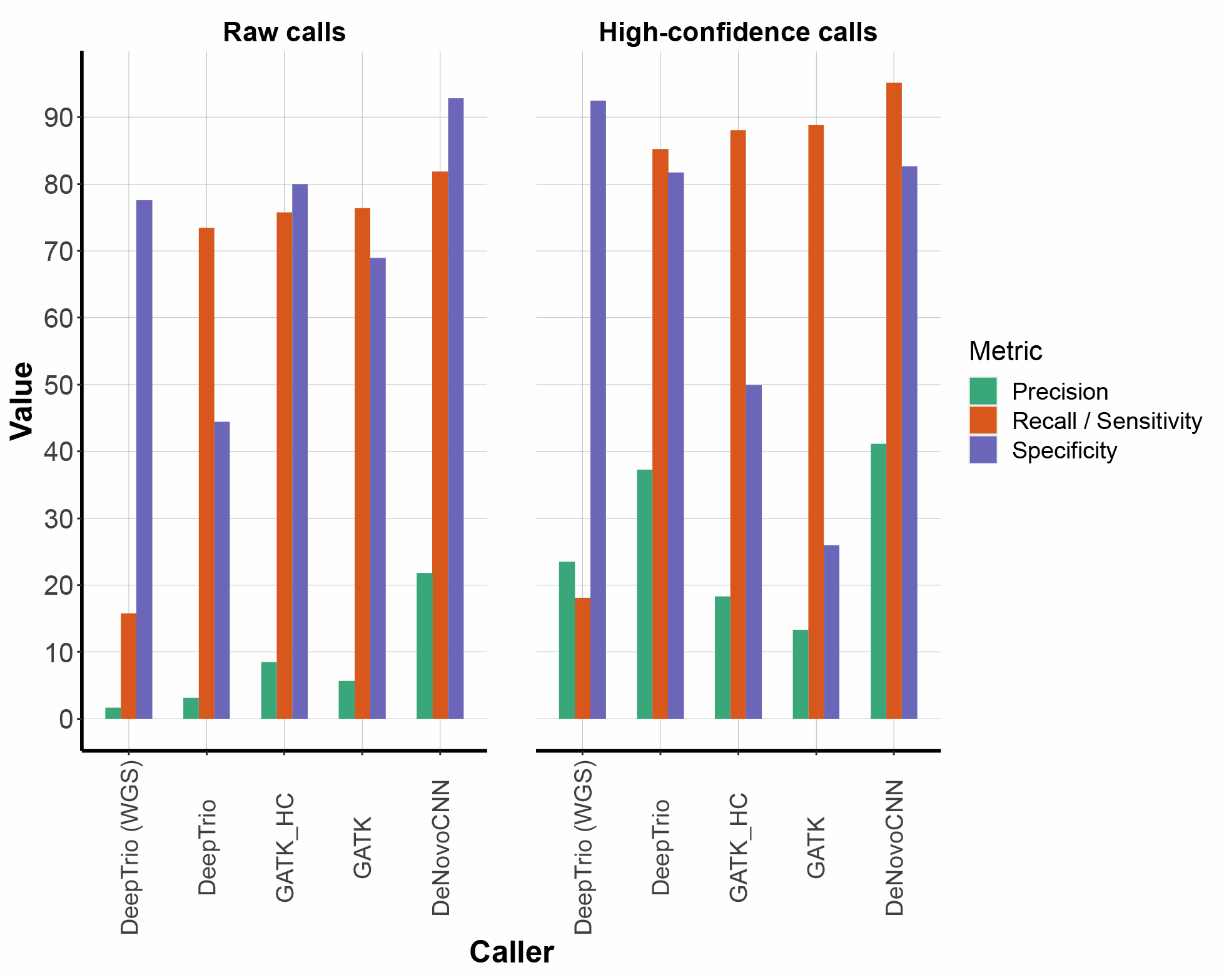
Validation and comparison of the DeNovoCNN method on genome data. The figure contains 2 plots for the performance based on raw and high confidence DNM calls (Material and methods). The tools used for the comparison are on the horizontal axis: DeepTrio (DeepVariantWGS), DeepTrio (DeepVariant_unfiltered), GATK with high quality DNMs, GATK, and DeNovoCNN. The green, orange and violet bars show precision, recall (sensitivity), and specificity respectively.

**Table 3.**
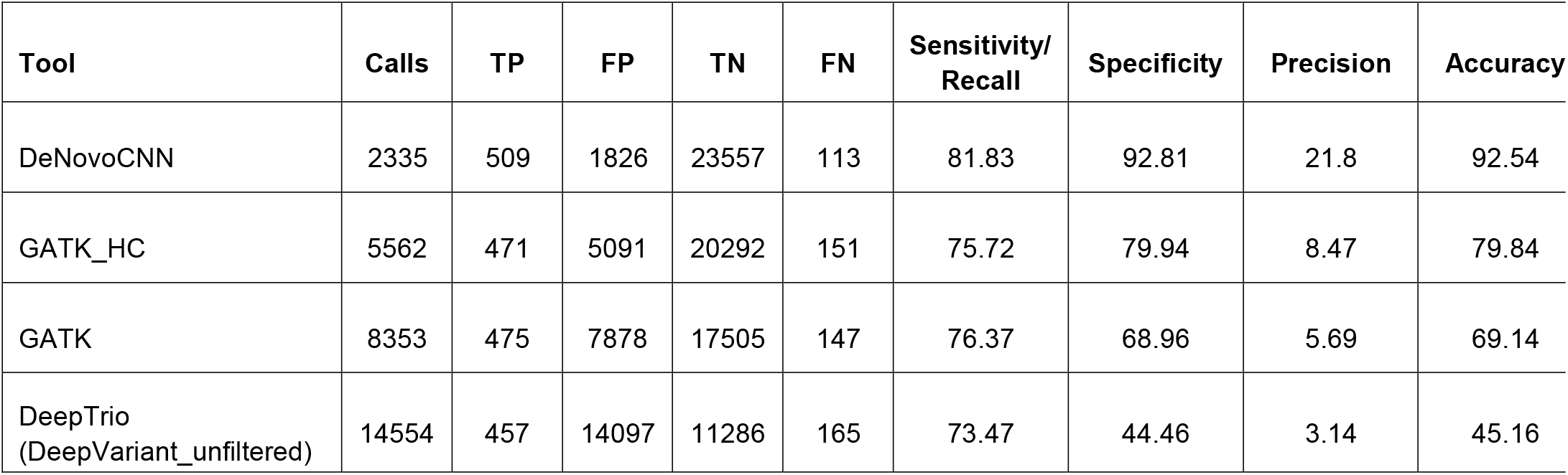
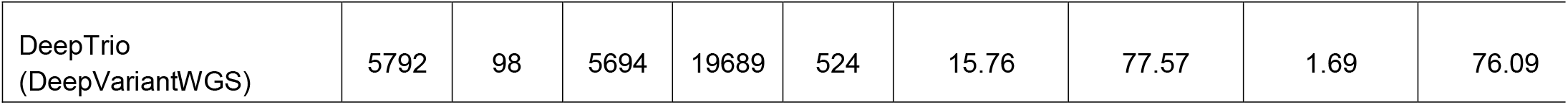
Validations on the 7 WGS trios using PacBio LRS based on raw calls of the tools. Every row shows different statistics and performance metrics for DeNovoCNN, GATK with high quality DNMs and GATK, and DeepTrio with two different settings. The first column shows the name of the tool, the next five columns show the number of total DNM calls, the number of true positive, false positive, true negative and false negative DNM calls respectively based on comparison with DNMs from PacBio HiFi LRS. The next 4 columns show the performance metrics, such as sensitivity/recall, specificity, precision and accuracy.

| Tool | Calls | TP | FP | TN | FN | Sensitivity/<br>Recall | Specificity | Precision | Accuracy |
| --- | --- | --- | --- | --- | --- | --- | --- | --- | --- |
| DeNovoCNN | 2335 | 509 | 1826 | 23557 | 113 | 81.83 | 92.81 | 21.8 | 92.54 |
| GATK_HC | 5562 | 471 | 5091 | 20292 | 151 | 75.72 | 79.94 | 8.47 | 79.84 |
| GATK | 8353 | 475 | 7878 | 17505 | 147 | 76.37 | 68.96 | 5.69 | 69.14 |
| DeepTrio<br>(DeepVariant_unfiltered) | 14554 | 457 | 14097 | 11286 | 165 | 73.47 | 44.46 | 3.14 | 45.16 |
| DeepTrio<br>(DeepVariantWGS) | 5792 | 98 | 5694 | 19689 | 524 | 15.76 | 77.57 | 1.69 | 76.09 |

## DISCUSSION

Here we introduced DeNovoCNN, a novel approach to *de novo* variant calling based on a convolutional neural network. We applied DeNovoCNN to several independent datasets such as the GIAB WGS Ashkenazi trio, 20 WES trios and 7 whole-genome short-read sequencing trios to compare the performance with other methods (GATK, Samtools, DeepTrio, DeNovoGear) based on orthogonal validations with the GIAB gold standard DNM dataset, Sanger/IonTorrent validations and PacBio LRS respectively. For all of these datasets, we find that DeNovoCNN consistently outperforms other approaches in terms of precision, recall and specificity. An advantage of DeNovoCNN is that it is in principle not dependent on the variant calling method itself, which will make it easier to use on existing datasets and incorporate into existing data analysis pipelines. Other approaches such as GATK and DeepTrio, require that variants are called in a specific way and otherwise will not be able to produce DNM calls. This means that for existing datasets recalling is likely to be needed which may represent a significant (computational) effort. However, we cannot fully exclude that for particular variant calling methods, DeNovoCNN will need to be retrained.

Another potential advantage comes from the observation that DeNovoCNN seems to be able to identify also mosaic variants. Because these variants are even rarer than DNMs we were not able to perform a thorough evaluation. However, we ran DeNovoCNN on 10 WES trios in which a mosaic variant had been reported and identified 9 out of 10 of these variants (Supplementary Table 11).

The biggest challenge of DeNovoCNN is common for most deep learning models and is related to the fact that the model should generalize well to other data unseen during training. Deep learning models can be sensitive to subtle differences in the data which can lead to unexpected results. Because we exclusively used data from a single center for the training of our model, albeit from two different capture kits and sequencing instruments we have used data augmentation techniques to prevent overfitting. Our validations are for the most part done on completely different datasets than those used for training the model, such as the GIAB dataset, the 7 WGS trios, and the WES data from the Solve-RD project. For all of these datasets, we obtain good results, supporting the notion that DeNovoCNN is not overly biased towards the training dataset. However, we cannot exclude that some residual bias from the training data exists and that this may become apparent when DeNovoCNN is applied to other datasets. We tested for two potential biases, namely lower coverage sequencing data, and omitting Base Quality Score Recalibration (BQSR). At lower coverages, we observed no drastic effects on recall and precision (Supplementary Figure 10) and find a high correlation in DeNovoCNN predictions for the alignments with and without BQSR (Supplementary Figure 11). Another hurdle to the application of DeNovoCNN could be the fact that nowadays CRAM is becoming a new data standard (23) for storing sequence alignments. Therefore, we also compared DeNovoCNN results from BAM files and corresponding CRAM files with quality-score binning. We find that the correlation between the prediction on BAM and CRAM files is very high (Supplementary Figure 12) and therefore expect DeNovoCNN to work equally well on CRAM files.

A possible source of bias of our model lies in the generation of the training data using manual inspection of the variants in IGV. We did not observe any obvious biases in the performance of DeNovoCNN on independent datasets that seemed to arise from our manual inspection. However, comprehensive gold standard DNM datasets, including DNMs in the complex regions of the human genome, will be needed to make such manual visual inspection unnecessary in the future.

Although DeNovoCNN shows overall good performance compared to other methods, we have seen some possibilities for future improvements. We noticed that the performance on indels is lower than the performance on substitutions. This is not surprising since the calling of indels is also more challenging than for substitution variants, but could also be explained by the fact that the amount of the indels events in the training dataset is much lower than substitutions events. In addition, DNMs were mostly validated using visual inspection. Because indel *de novo* events are more difficult to distinguish manually from sequencing artefacts, this could have led to a poorer training dataset specifically for these events. In general, we remark that for future improvements in DNM detection, it will be essential to have sizeable, curated training datasets.

Other possible improvements could lie in the model itself. The training of deep learning models is computationally intensive, which is why we chose a relatively simple CNN. Although we did not observe any clear negative effects on model performance, more complex architectures and more context information for variants could potentially improve the performance further.

## Supporting information

GIAB.manual.validation

20_WES_comarison_FP

## AVAILABILITY

DeNovoCNN is open-source software available as a Docker container in the GitHub repository (https://github.com/Genome-Bioinformatics-RadboudUMC/DeNovoCNN).

The training dataset for DeNovoCNN is available in the GitHub repository (https://github.com/Genome-Bioinformatics-RadboudUMC/DeNovoCNN_training_dataset).

## ACKNOWLEDGEMENT

The aims of this study contribute to the Solve-RD project (to CG and LELMV) which has received funding from the European Union’s Horizon 2020 research and innovation programme under grant agreement No 779257.

## FUNDING

This project was partly funded by The Netherlands Organisation for Scientific Research (917-17-353) to CG.

## CONFLICT OF INTEREST

The authors declare that there is no conflict of interest.

## Notes

https://github.com/Genome-Bioinformatics-RadboudUMC/DeNovoCNN

https://github.com/Genome-Bioinformatics-RadboudUMC/DeNovoCNN_training_dataset

### Competing Interest Statement

The authors have declared no competing interest.

