## Supplementary Data for "DeNovoCNN: A deep learning approach to *de novo* variant calling in next generation sequencing data"

### SUPPLEMENTARY METHODS

#### GIAB WGS dataset

In order to compare the performance of DeNovoCNN with existing tools, we decided to use BAM and VCF files (Supplementary links) of the extensively validated Ashkenazim Trio (NA24385; NA24149; NA24143) from the Genome in a Bottle (GIAB) consortium (1). BAM files base recalibration was performed with GATK V.3.4-46. This trio was whole-genome sequenced using various different technologies in order to create a complete dataset of the high-quality cell line or germline 1323 DNMs that are Mendelian inconsistent variants (Supplementary links) filtered in a way that the son was heterozygous and both parents were a homozygous reference.

We compared our results to GATK, GATK filtered for high confidence DNM calls (GATK_HC), DeNovoGear filtered using 0.5 and 0.9 probability thresholds (DeNovoGear-0.5, DeNovoGear-0.9) and DeepTrio, applied on BAMs with and without BQSR and using DeepTrio_WGS and DeepTrio_unfiltered configs for GLnexus (Supplementary Figure 5).

For 74 variants (38 substitutions, 10 insertions and 26 deletions) that are part of the GIAB high quality DNM list, there were no DNM calls by any of the evaluated tools. Upon visual inspection in IGV, some of these variants appear to be inherited or are located in a highly repetitive region which might have an effect on the performance metrics. We performed a manual examination of the whole set of high quality DNM independently by two specialists labelling them as *de novo,* not *de novo* and unknowns, and left only those that were confirmed as DNM by both specialists resulting in 1106 DNMs (Supplementary File GIAB.manual.validation.xls).

#### In-house 20 WES trios dataset

In order to compare the DeNovoCNN performance to existing methods for identifying DNMs we selected 20 internal WES samples that were not used during training and validation. These samples were exome sequenced on an Illumina HiSeq4000 using the Agilent SureSelect v5 exome kit. Medium target coverage was on average 110x. BAM files were generated using BWAmem V.0.7.13 and then a base recalibration was performed with GATK v.3.4-46. For CRAM files bamUtil v.1.0.14 was used for quality scores binning and Samtools v.1.10 was used for BAM to CRAM conversion. VCF files were generated using GATK v.3.4-46, but no joint calling was performed.

We used our in-house *de novo* tool, GATK, DeNovoGear-0.5 and DeepTrio as commonly used *de novo* calling algorithms and performed an in-depth comparison of all individual DNM calls between the five algorithms. DeepTrio was applied on BAMs with and without BQSR and using DeepTrio_WES and DeepTrio_unfiltered configs for GLnexus. For all the tools variants were only called in the coding regions

according to Gencode v.31 for GRCh37 with an extension of 20bp intronic sequence at both sides. We combined all DNMs that were called by at least one of the tools and manually inspected them in IGV, discarding the variants that are obviously not *de novo* (e.g. were clearly inherited from the parents or located in a difficult region with a lot of sequencing mistakes around). For the remaining 50 variants, we performed standard Sanger/IonTorrent sequencing to confirm whether they truly were *de novo.*
We compare the tools based on raw calls as well as using postprocessing of the calls described in Materials and Methods.

**Multi-platform trio dataset**

Because the training of the model was done on in-house samples, we wanted to exclude the possibility that our model was biased towards our particular type of sequencing and exome enrichment method. Therefore, we performed a more extensive evaluation using 551 trios from the SolveRD consortium, where the number of *de novo* calls from GATK and DeNovoCNN doesn't exceed 100. The SolveRD consortium contains WES trio data from a variety of different sequencing platforms and enrichment kits (Supplementary Table 7). Mapping and calling were performed using the GPAP platform (2,3) with BWAmem v.0.7.8 for BAM file generation and GATK v.3.6-0 for gVCF files generation (4,5). Further gVCF to VCF conversion was done using bcftools v.1.9. Only variants with GATK quality score >100 were included. For CRAM files bamUtil v.1.0.14 was used for quality score binning and Samtools v.1.10 was used for BAM to CRAM conversion.

**WGS dataset**

*Long read sequencing*

We sequenced 7 trios on the Pacific Biosciences Sequel II instrument with 3 SMRT cells per sample, targeting a 30x coverage with HiFi reads. Consensus HiFi reads were generated with ccs 4.2.0, that was followed by with https://github.com/williamrowell/pbRUGD-workflow/ processing pipeline.

We then aligned those reads to the GRCh38/Hg38 genome with pbmm2 v.1.4.0, using default parameters. For SNV calling, we ran DeepVariant v. 1.1.0 (6) with default setting.

We filtered de novo mutations using slivar v.0.2.7 (7). High quality *de novo* mutations list was generated using the following filters: parental genotype is 0/0, proband genotype is 0/1, parental alternative allele depth is 0, proband allele depth $>5$, reference allele depth $\geq 10$, total depth $\leq 50$, quality score $\geq 30$, genotype score $\geq 20$ and allele count $<5$ in gnomAD and HPRC controls.

*Short read sequencing*

7 WGS trios were sequenced at 50x coverage on Illumina NovaSeq 6000 instruments. BAM files were generated using BWA V.0.78. For CRAM files bamUtil v.1.0.14 was used for quality scores binning and Samtools v.1.10 was used for BAM to CRAM conversion. VCF files were generated using GATK v.3.8

We applied DeNovoCNN, GATK and DeepTrio on short read sequencing data described above. For GATK *de novo* calls were generated after recalling using GATK best practices pipeline for germline short variant discovery (SNPs + indels) and “Genotype Refinement workflow for germline short variants”. DeepTrio was applied using two different settings for GLnexus: with no filtering applied (DeepVariant_unfiltered) and with optimized filtering settings for WGS (DeepVariant_WGS). *De novo* calls from the tools were compared with the set of the high quality *de novo* calls obtained from long read sequencing.

#### BAM and CRAM dataset

To show that DeNovoCNN predictions are stable for cramming procedure a dataset of 3 WES trios was used. Data was sequenced at 180x medium coverage on Illumina NovaSeq 6000 using Twist Human RefSeq Panel. BAM and VCF files were generated the same way as for 20 WES in-house dataset. For CRAM files bamUtil v.1.0.14 was used for quality scores binning and Samtools v.1.10 was used for BAM to CRAM conversion. We applied DeNovoCNN twice on these data, first using BAM files as an input, and second using CRAM files as an input, and compared the predicted probabilities on all the variants.

### SUPPLEMENTARY LINKS

#### GIAB Ashkenazim Trio data

- Son VCF ftp://ftp-trace.ncbi.nlm.nih.gov/giab/ftp/release/AshkenazimTrio/HG002_NA24385_son/NISTv3.3.2/GRCh37/HG002_GRCh37_GIAB_highconf_CG-IllFB-IllGATKHC-Ion-10X-SOLID_CHROM1-22_v.3.3.2_highconf_triophased.vcf.gz + .tbi file
- Father VCF ftp://ftp-trace.ncbi.nlm.nih.gov/giab/ftp/release/AshkenazimTrio/HG003_NA24149_father/NISTv3.3.2/GRCh37/HG003_GRCh37_GIAB_highconf_CG-IllFB-IllGATKHC-Ion-10X_CHROM1-22_v.3.3.2_highconf.vcf.gz + .tbi file
- Mother VCF ftp://ftp-trace.ncbi.nlm.nih.gov/giab/ftp/release/AshkenazimTrio/HG004_NA24143_mother/NISTv3.3.2/GRCh37/HG004_GRCh37_GIAB_highconf_CG-IllFB-IllGATKHC-Ion-10X_CHROM1-22_v.3.3.2_highconf.vcf.gz + .tbi file
- Son BAM ftp://ftp.ncbi.nlm.nih.gov/giab/ftp/data/AshkenazimTrio/HG002_NA24385_son/NIST_HiSeq_HG002_Homogeneity-10953946/HG002Run01-11419412/HG002run1_S1.bam

- Father BAM ftp://ftp.ncbi.nlm.nih.gov/giab/ftp/data/AshkenazimTrio/HG003_NA24149_father/NIST_HiSeq_HG003_Homogeneity-12389378/HG003Run01-13262252/HG003Run01_S1.bam
- Mother BAM ftp://ftp.ncbi.nlm.nih.gov/giab/ftp/data/AshkenazimTrio/HG004_NA24143_mother/NIST_HiSeq_HG004_Homogeneity-14572558/HG004Run01-15133132/HG004Run01_S1.bam
- Mendelian inconsistent variants ftp://ftp-trace.ncbi.nlm.nih.gov/giab/ftp/release/AshkenazimTrio/HG002_NA24385_son/NISTv3.3.2/GRCh37/supplementaryFiles/HG002_GRCh37_GIAB_highconf_CG-IllFB-IllGATKHC-Ion-10X-SOLID_CHROM1-22_v.3.3.2_highconf_trioinconsistent.vcf.gz + tbi file
- High-quality regions ftp://ftp-trace.ncbi.nlm.nih.gov/giab/ftp/release/AshkenazimTrio/HG002_NA24385_son/NISTv3.3.2/GRCh37/HG002_GRCh37_GIAB_highconf_CG-IllFB-IllGATKHC-Ion-10X-SOLID_CHROM1-22_v.3.3.2_highconf_noinconsistent.bed

### SUPPLEMENTARY FIGURES

**Supplementary Figure 1. Overview of the dataset construction used for training, validation and test of the DeNovoCNN model.** The original 6,067 trios with 13,068 DNMs were manually divided in IGV into 10,274 DNMs, 1,174 inherited and 1,620 uncertain variants. Uncertain variants were discarded resulting in 5,616 trios. Existing IVs in these trios were randomly sampled leading to 55,134 IVs in total. Using this dataset, the interim DeNovoCNN model was trained. To improve the representativity of difficult regions for DNM calling, 403 trios were randomly chosen to apply the interim DeNovoCNN model. After manual inspection of the calls in IGV 159 DNMs and 905 IVs were added to the dataset. The resulting dataset was divided into train, validation and test datasets using a 70/15/15 per cent ratio. Two artificial trios with unrelated parents were added to the training dataset to increase the number of DNMs by 1,005.

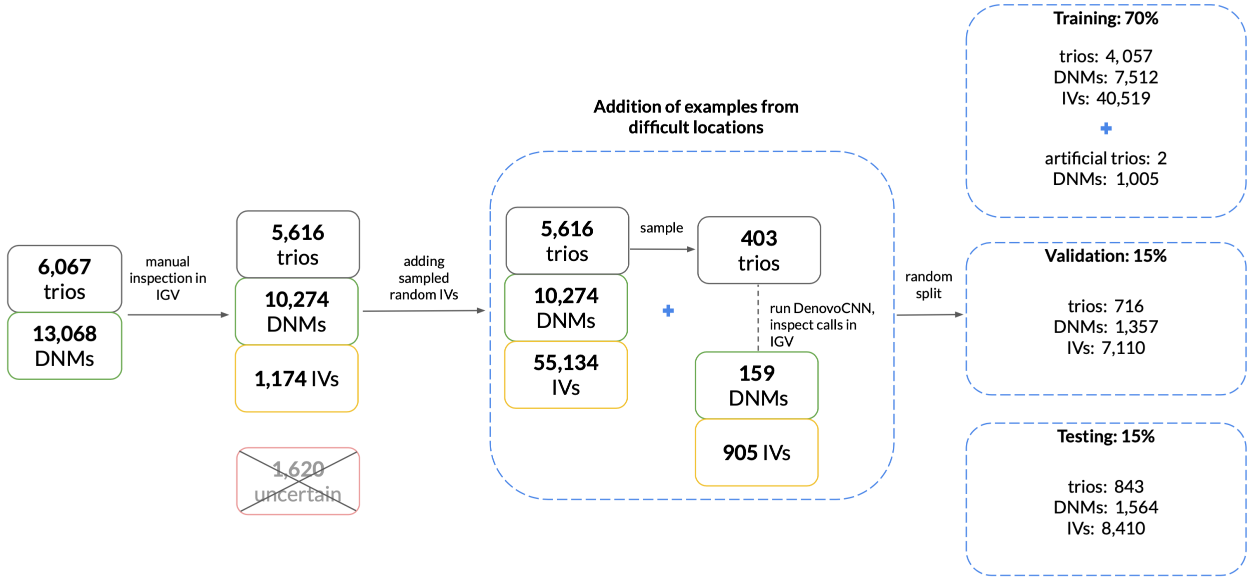

**Supplementary Figure 2. Convolutional neural network architecture.** The CNN architecture is shown from left to the right. Each yellow block shows a 3x3 convolution layer, each light blue block depicts a batch normalization layer, green blocks correspond to the squeeze-and-excitation layer, and in the grey block, the concatenation of global maximum and average pooling is shown. The green sphere shows the sigmoid function that gives the final probability of the variant being *de novo.*

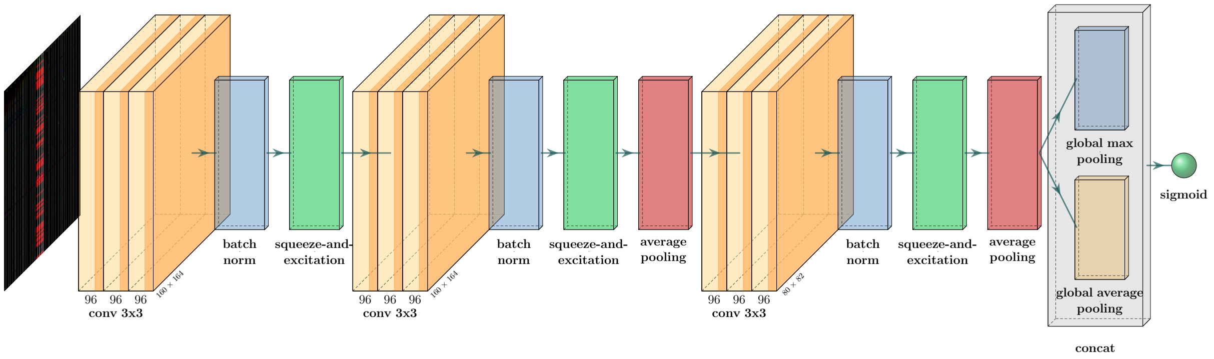

**Supplementary Figure 3. Examples of RGB-image-like representations of a *de novo* substitution, deletion and insertion based on sequence alignment data.** The central part of the image marks the region of interest and has higher pixel intensity. The red colour channel represents bases found in the child, and the green and blue channels represent the parents. Columns correspond to one-hot encoded bases, while rows correspond to different reads.

A

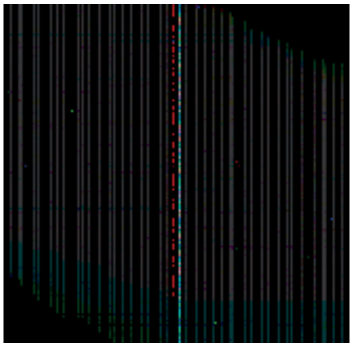

De novo substitution

B

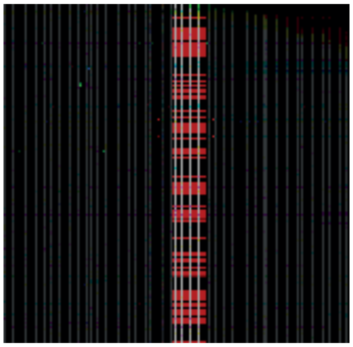

De novo deletion

C

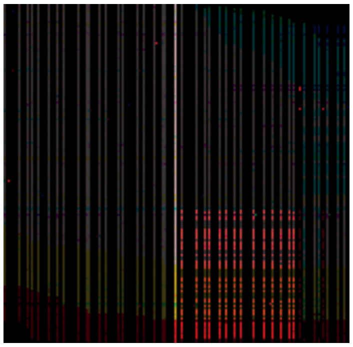

De novo insertion

**Supplementary Figure 4. Learning curves from training DeNovoCNN.** The figures show the performance in terms of AUC (on the left y-axis) and binary cross-entropy (on the right y-axis) of the neural networks at the end of each epoch during training. Red and blue colours correspond to the training and validation datasets respectively. The marks at the plots show the model that was automatically selected as the final based on the performance on the validation dataset.

A. Substitutions

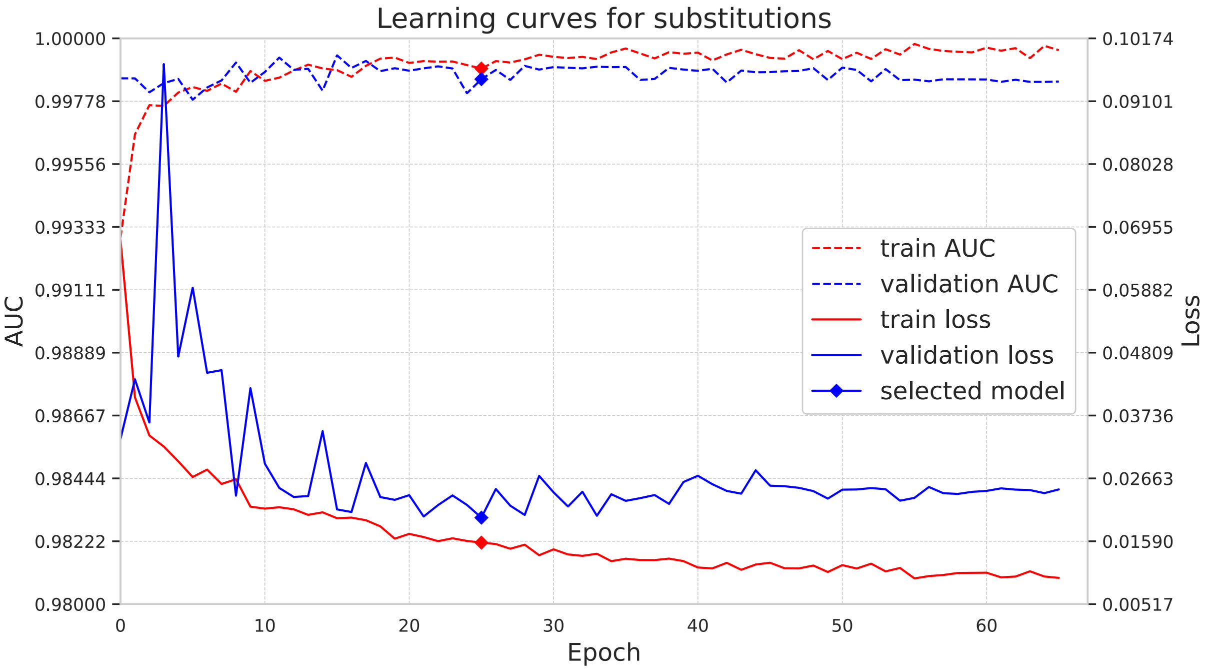

B. Deletions

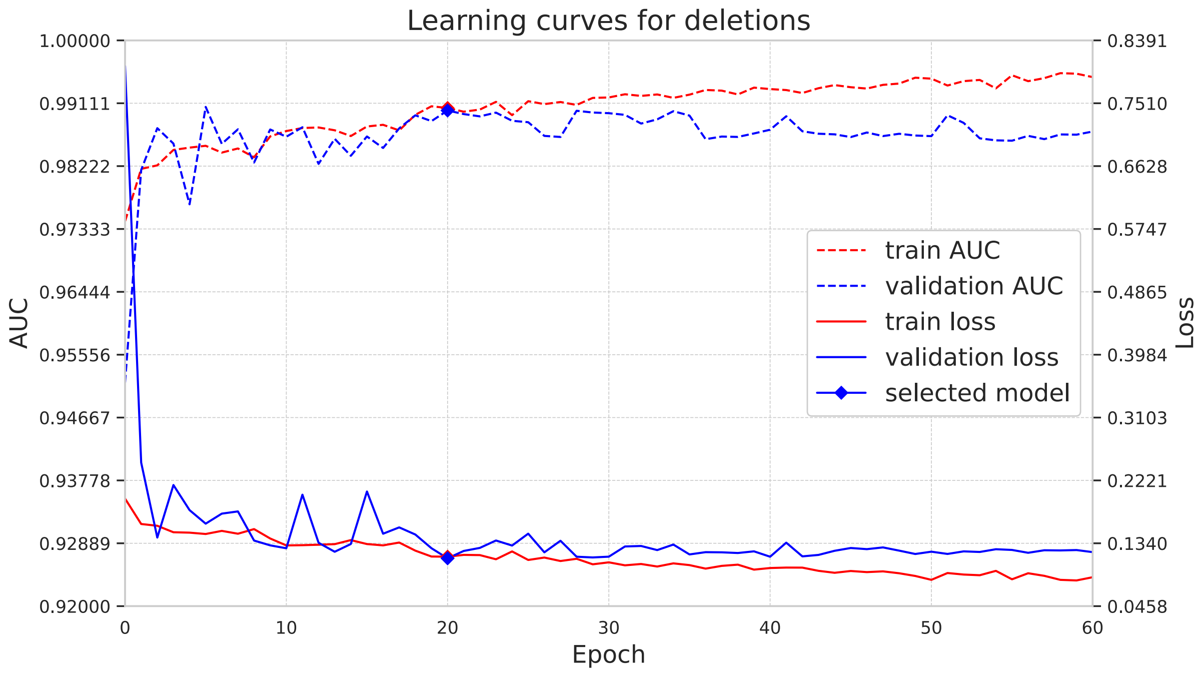

C. Insertions

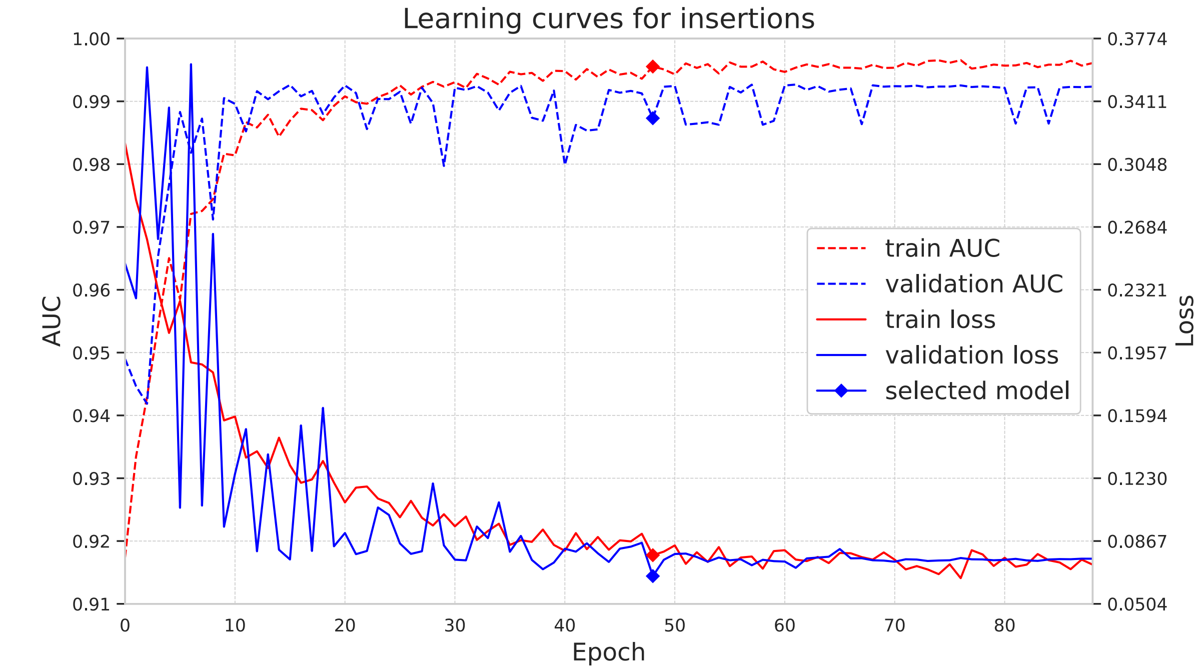

**Supplementary Figure 5. DeNovoCNN results on the test dataset.** The red line shows DeNovoCNN performance on substitutions, the blue line shows the performance on deletions and the green line on insertions. Each line shows precision and recall values for all possible thresholds on DNM probability values. The recall is on the horizontal axis; the vertical axis corresponds to precision. Average precision (AP/PR-AUC) summarizes the graphs as the area under the precision-recall curve.

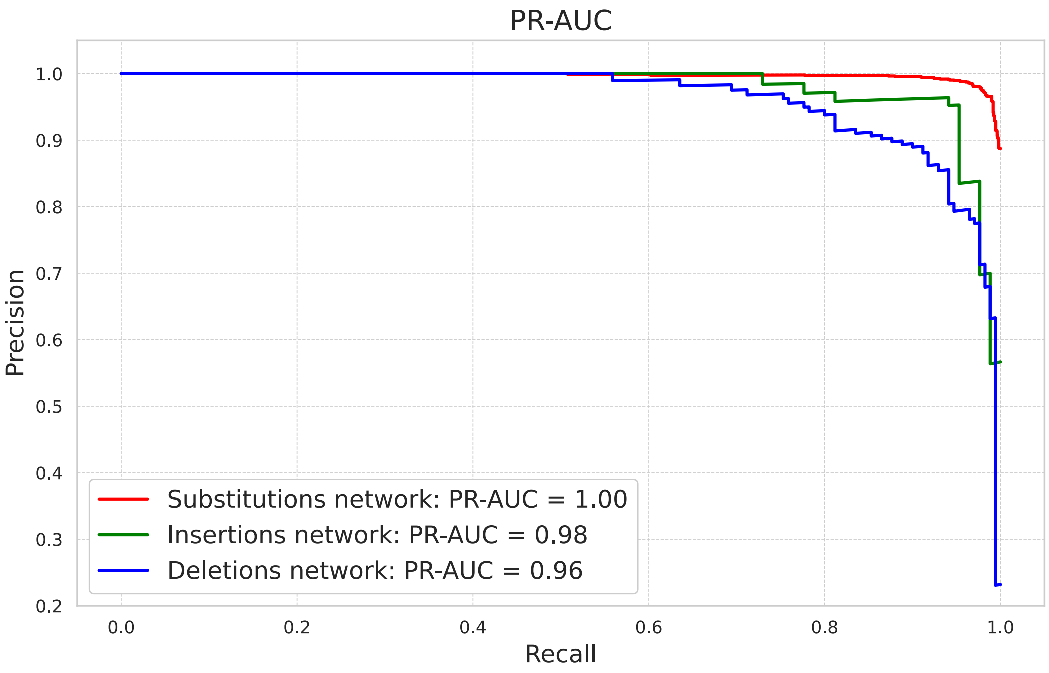

**Supplementary Figure 6. Feature maps from the last convolution layer of DeNovoCNN indicate features that are used by the model.** Red areas show the regions of the most interest for CNN. It is shown that the network uses both information about the context and the variation itself. Subplots A-C were generated using the substitutions model, showing interest in (A) the *de novo* variant area, (B) the context of the variant, (C) coverage information; subplots D-F were generated using the insertions model, showing interest in (D) *de novo* variant area, (E) context after a variant of interest, (F) context before a variant of interest; subplots G-I were generated using deletions model, showing interest in (G) *de novo* variant area, (H) context of the variant, (I) coverage information.

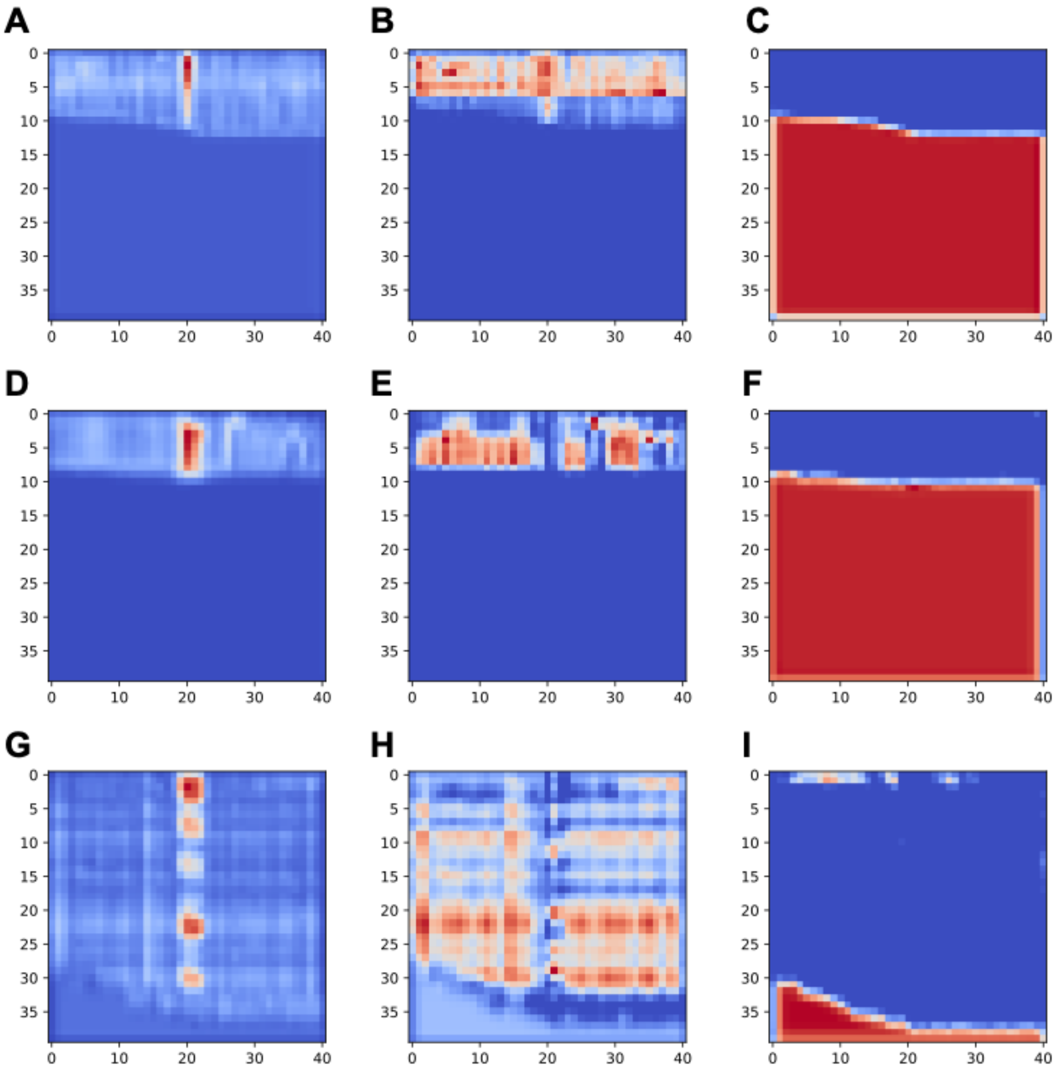

**Supplementary Figure 7. DeNovoGear probabilities distribution on GIAB data.** The plot shows the distribution of DNM probability values ≥0.5 predicted by DeNovoGear on the GIAB dataset. Based on the distribution of probabilities we choose a cutoff of 0.5 and 0.9, since the vast majority of de novo calls lies in the area of the probability ≥0.95.

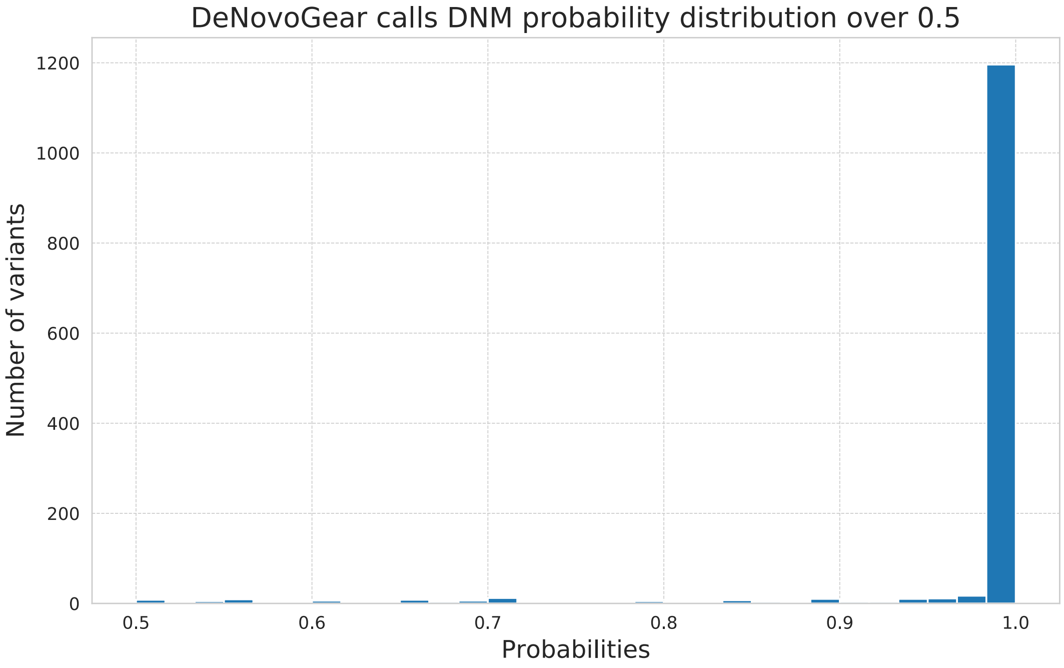

**Supplementary Figure 8. Validation process on 20 WES trios.** DeNovoCNN, DeNovoGear, GATK, and our in-house tool were applied to our in-house 20 WES trios dataset. All calls then were manually validated in IGV to create a dataset of 50 candidate variants that look like potential *de novo*. The set of 50 variants then were sent to additional Sanger/IonTorrent sequencing to confirm 24 variants as true *de novo*

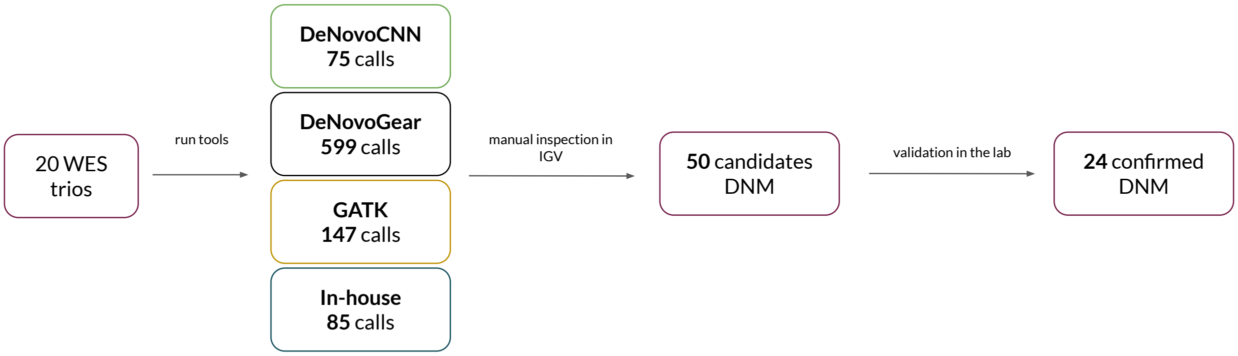

.

**Supplementary Figure 9. Violin plots of the distribution of the number of DNMs per sample, grouped per enrichment kit/sequencing combination.** The plot shows the smoothed distribution of the number of trio DNM calls per enrichment kit. The numbers between parentheses are the number of samples for the corresponding sequencing platform.

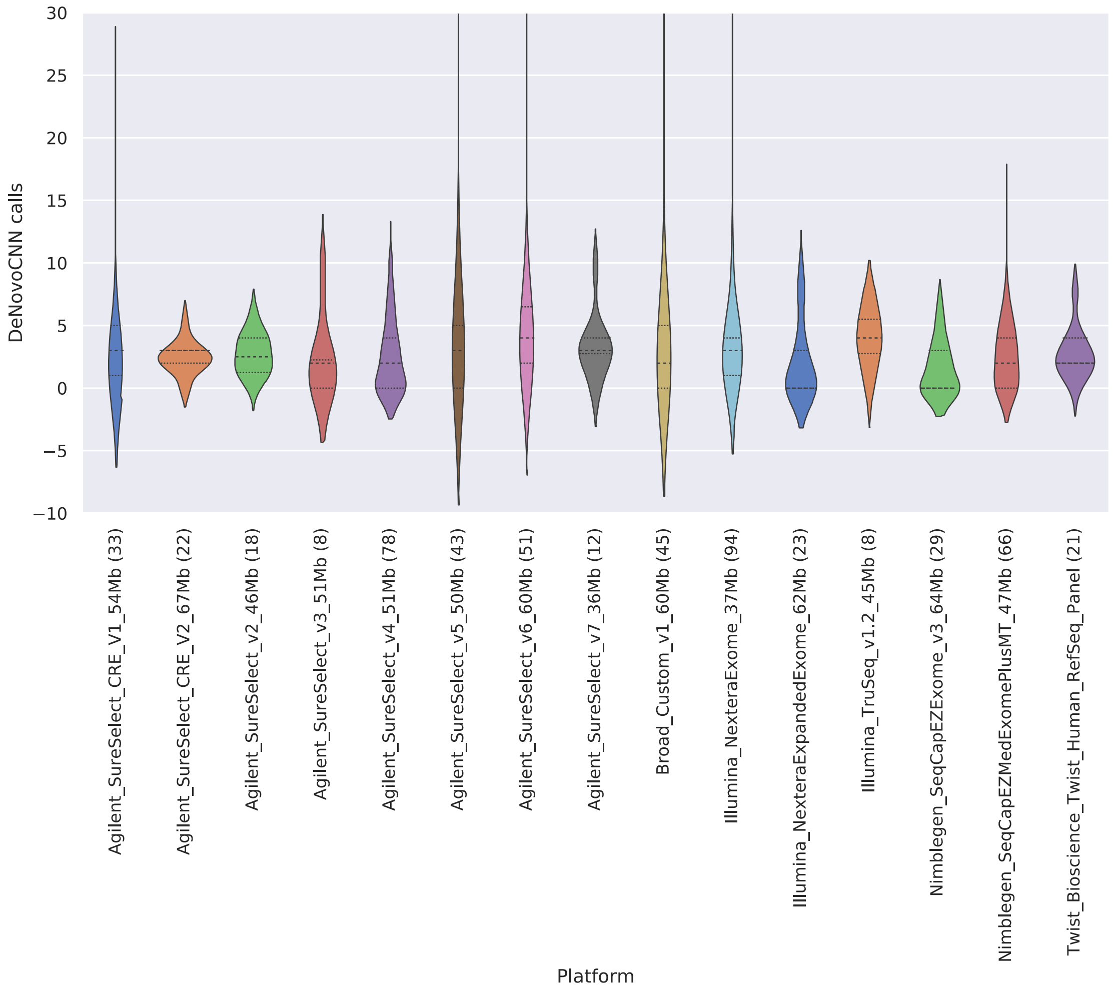

**Supplementary Figure 10. DeNovoCNN DNM performance on different coverages.** The plot shows the change in recall (left plot, y-axis) and precision (right plot, y-axis) with the decrease of the sequencing coverage (x-axis).

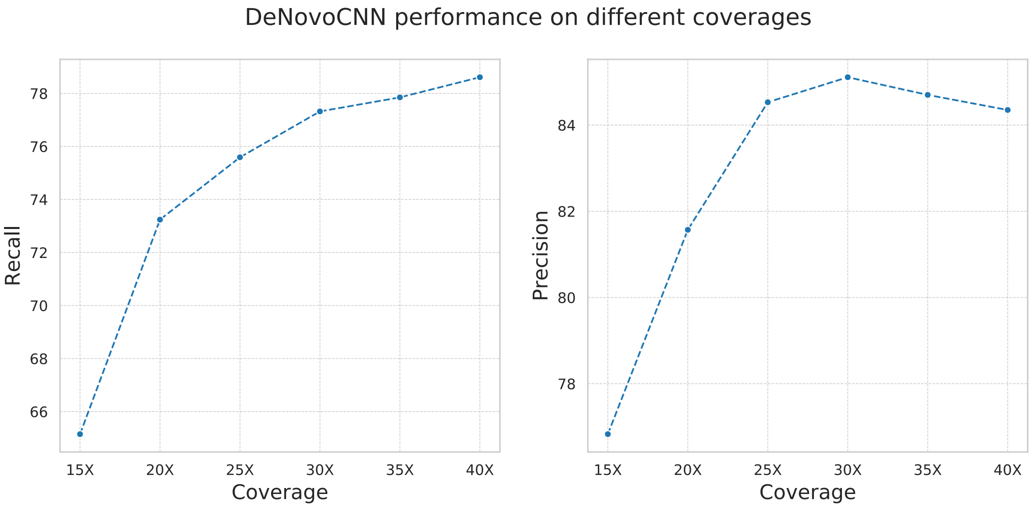

**Supplementary Figure 11. DeNovoCNN DNM probabilities scatter plot on BAM with and without BQSR on 20 WES trios.** Scatter plot of DNM probabilities predicted by DeNovoCNN based on BAM with BQSR (on the horizontal axis) and BAM without BQSR (on the vertical axis) for 20 WES trios. The results show that predictions are highly correlated (correlation coefficient of 0.9916).

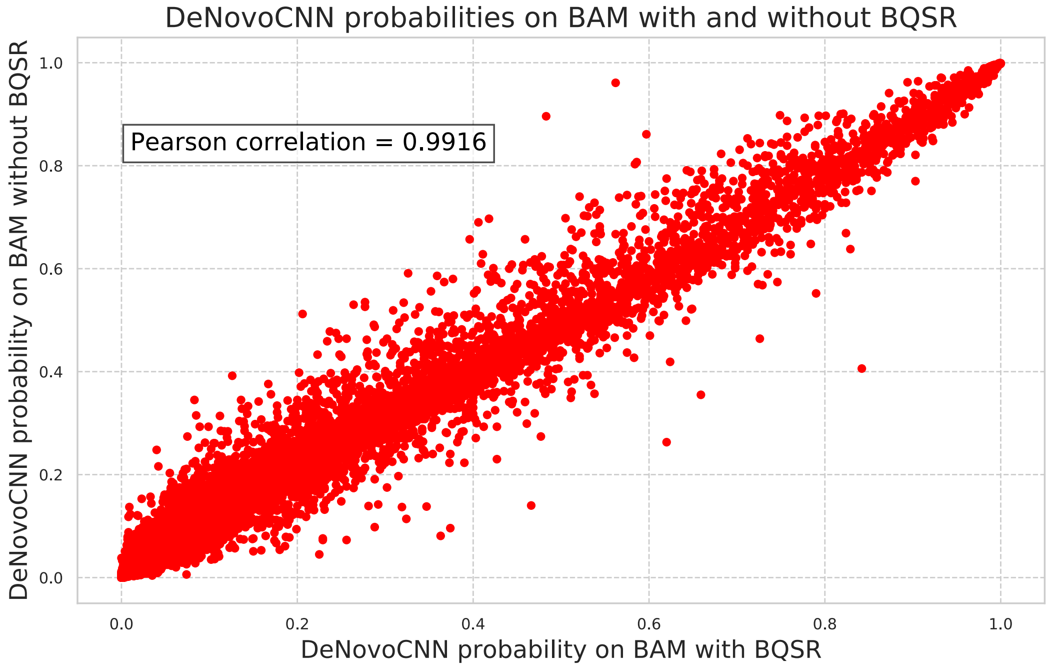

**Supplementary Figure 12. DeNovoCNN DNM probabilities scatter plot on BAM and CRAM data.** Scatter plot of DNM probabilities predicted by DeNovoCNN based on BAM (on the horizontal axis) and CRAM (on the vertical axis) files. The results show that predictions are highly correlated (correlation coefficient of 0.9886).

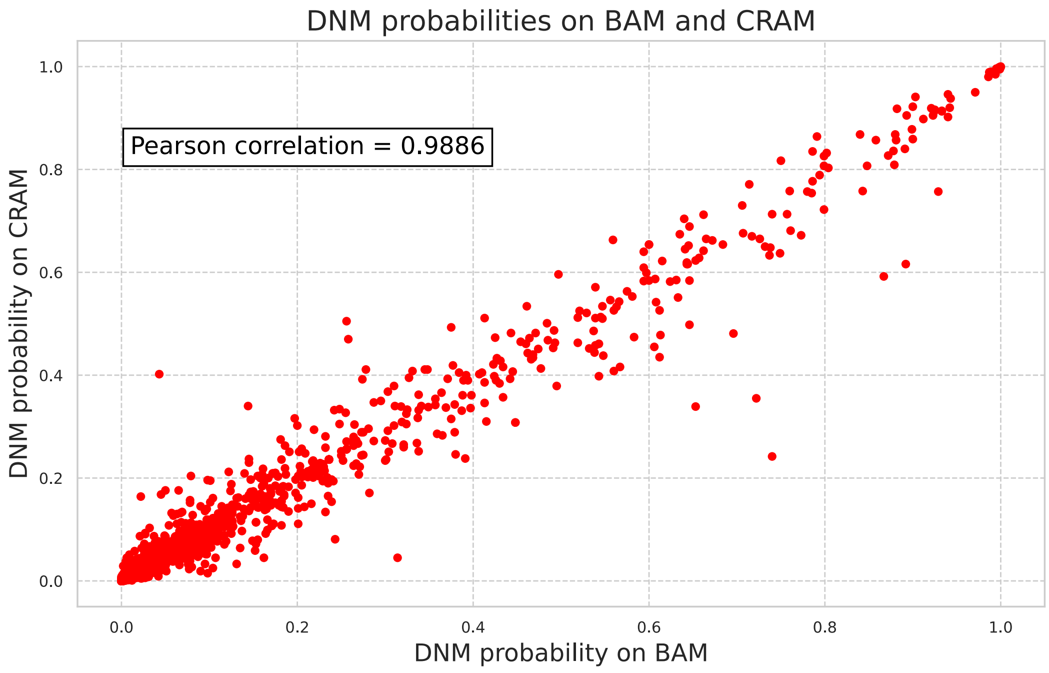

### SUPPLEMENTARY TABLES

**Supplementary Table 1. Overview of the training, validation and test datasets.** The table shows the number of variants for training, validation and test datasets for each class: *de novo* mutations (DNM) and inherited variants (IV). Every row shows different types of variants: substitutions, deletions, insertions and all types of variants together (total). The last line shows the number of trios for each dataset.

| **Dataset** | **Training** | | **Validation** | | **Testing** | |
|---|---|---|---|---|---|---|
| | DNM | IV | DNM | IV | DNM | IV |
| **Substitutions** | 6,179 | 35,488 | 1,126 | 6,186 | 1,309 | 7,322 |
| **Deletions** | 1,350 | 2,732 | 156 | 482 | 170 | 594 |
| **Insertions** | 988 | 2,410 | 75 | 442 | 85 | 494 |
| **Total** | 8,517 | 40,590 | 1,357 | 7,110 | 1,564 | 8,410 |
| **# of trios** | 4,057 + 2 artificial | | 716 | | 843 | |

**Supplementary Table 2. Hyperparameters for the training of neural networks.** Each row contains the information of one hyperparameter of the network, such as the value for the substitutions, deletions and insertions networks respectively, and the optimization space used for the hyperparameter search.

| **Parameter** | **Substitution network** | **Deletion network** | **Insertion network** | **Optimization space** |
|---|---|---|---|---|
| Convolution kernel size | 3x3 | 3x3 | 3x3 | - |
| Number of convolutional features | 96 | 96 | 96 | [32, 64, 96, 128] |
| L1 regularization coefficient of the last layer | 3.1e-7 | 3.1e-7 | 3.1e-7 | [1e-10, 0.1] |
| Batch size | 64 | 64 | 32 | [32, 64] |
| Learning rate | 0.00046 | 0.0025 | 0.0026 | [1e-8, 0.01] |
| Learning rate decay factor | 0.5 | 0.5 | 0.5 | - |
| Learning rate decay step (in epochs) | 10 | 10 | 10 | - |
| Adam weight decay | - | 1.1e-6 | 7.3e-8 | [1e-8, 0.01] |
| Maximum number of epochs | 100 | 100 | 100 | - |
| Early stop patience | 40 | 40 | 40 | - |

**Supplementary Table 3. Overview of datasets used in training, validation and testing of DeNovoCNN.** Each row shows the description (type of data, purpose of usage, number of trios and source) of every dataset used in training, validation and testing of DeNovoCNN.

| **Dataset** | **Purpose** | **# Trios** | **Source** |
|---|---|---|---|
| WES, 110x coverage, Agilent SureSelect v4 and v5 | DeNovoCNN training dataset | 4,057 + 2 artificial | Radboudumc |
| WES, 110x coverage, Agilent SureSelect v4 and v5 | DeNovoCNN validation dataset, hyperparameter optimization | 716 | Radboudumc |
| WES, 110x coverage, Agilent SureSelect v4 and v5 | DeNovoCNN test dataset, performance assessment | 843 | Radboudumc |
| WES, 110x coverage, Agilent SureSelect v5 | Comparison of DNM callers on WES | 20 | Radboudumc |
| WGS, 50x coverage on Illumina NovaSeq 600 + 30x coverage Pacific Biosciences Sequel II | Comparison of DNM callers on WGS | 7 | Radboudumc |
| WGS | Comparison of DNM callers on external WGS data | 1 | GIAB |
| WES | DeNovoCNN validation on different sequencing platforms and kits | 551 | Solve-RD |
| WES, 180x coverage, Twist Human RefSeq Panel | Validation on BAM/CRAM | 3 | Radboudumc |

**Supplementary Table 4. Tools statistics on 20 WES dataset.** Each row shows the amount of de novo calls for the tools at every stage of the de novo variants filtering process for 20 WES trios. First row shows the statistics for the raw calling data, the second row shows only variants, that are within the target bed file, the next row shows the data for exonic variants only, the fourth row shows the result of the variants intersection with the ones from GATK HaplotypeCaller, and the last row shows the result of filtering DNMs based on in-house filters (Materials and methods).

| | **DeepTrio (DeepVariant_unfiltered, BQSR)** | **DeepTrio (DeepVariant_unfiltered, no BQSR)** | **DeepTrio (DeepVariantWES, BQSR)** | **DeepTrio (DeepVariantWES, no BQSR)** | **GATK** | **DeNovoGear** | **DeNovoCNN** |
|---|---|---|---|---|---|---|---|
| Raw calls | 38,449 | 38,531 | 20,558 | 20,530 | 9,352 | 1,566 | 1,650 |
| Within target file | 4,239 | 4,241 | 1,683 | 1,675 | 353 | 694 | 121 |
| Exonic variants | 3,348 | 3,364 | 1,290 | 1,275 | 147 | 599 | 75 |
| Intersection with HaplotypeCaller | 2,527 | 2,539 | 1,086 | 1,081 | 134 | 398 | 75 |
| High quality variants | 1,457 | 1,490 | 685 | 672 | 38 | 84 | 30 |

**Supplementary Table 5. Performance of DeNovoCNN models on the test dataset.** Every row shows the performance metrics for the whole dataset and for substitution, deletion and insertion models respectively.

| Metric | Total | Substitutions | Insertions | Deletions |
|---|---|---|---|---|
| ROC AUC | 0.9988 | 0.9995 | 0.9957 | 0.9842 |
| Accuracy | 0.9895 | 0.9932 | 0.9827 | 0.9529 |
| Sensitivity/Recall | 0.9674 | 0.9771 | 0.9176 | 0.9176 |
| Specificity | 0.9936 | 0.996 | 0.9939 | 0.963 |
| F1 score | 0.9665 | 0.9775 | 0.9398 | 0.8966 |
| Precision | 0.9655 | 0.9778 | 0.963 | 0.8764 |
| True Positives | 1,513 | 1,279 | 78 | 156 |
| False Positives | 54 | 29 | 3 | 22 |
| True Negatives | 8,356 | 7,293 | 491 | 572 |
| False Negatives | 51 | 30 | 7 | 14 |

**Supplementary Table 6. Comparison on GIAB dataset.** Every row shows different statistics and performance metrics for DeNovoCNN, DeNovoGear with the probability threshold of 0.5 and 0.9, DeepTrio with different settings, GATK and high quality GATK, and our in-house tool. The first column shows the name of the tool, the next five columns show the number of total calls, the number of true positive, false positive, true negative and false negative calls respectively based on the GIAB reference DNMs. The next four columns show sensitivity (recall), specificity, precision and accuracy. All statistics are shown for substitutions (a), insertions (b), deletions (c), and all variants from chromosome 20 only (d).

a) Substitutions

| **Tool** | **Calls** | **TP** | **FP** | **TN** | **FN** | **Sensitivity/ Recall** | **Specificity** | **Precision** | **Accuracy** |
|---|---|---|---|---|---|---|---|---|---|
| DeNovoCNN | 1058 | 1046 | 12 | 455 | 64 | 94.23 | 97.43 | 98.87 | 95.18 |
| DeNovoGear-0.5 | 1324 | 1050 | 274 | 193 | 60 | 94.59 | 41.33 | 79.31 | 78.82 |
| DeNovoGear-0.9 | 1141 | 1034 | 107 | 360 | 76 | 93.15 | 77.09 | 90.62 | 88.4 |
| DeepTrio (DeepVariantWGS, BQSR) | 91 | 74 | 17 | 450 | 1036 | 6.67 | 96.36 | 81.32 | 33.23 |
| DeepTrio (DeepVariantWGS, no BQSR) | 165 | 151 | 14 | 453 | 959 | 13.6 | 97.0 | 91.52 | 38.3 |
| DeepTrio (DeepVariant_unfiltered, BQSR) | 1058 | 1014 | 44 | 423 | 96 | 91.35 | 90.58 | 95.84 | 91.12 |
| DeepTrio (DeepVariant_unfiltered, no BQSR) | 1080 | 1013 | 67 | 400 | 97 | 91.26 | 85.65 | 93.8 | 89.6 |
| GATK | 1112 | 1008 | 104 | 363 | 102 | 90.81 | 77.73 | 90.65 | 86.94 |
| GATK_HC | 1058 | 993 | 65 | 402 | 117 | 89.46 | 86.08 | 93.86 | 88.46 |
| In-house | 1076 | 1046 | 30 | 437 | 64 | 94.23 | 93.58 | 97.21 | 94.04 |

b) Insertions

| Tool | Calls | TP | FP | TN | FN | Sensitivity/ Recall | Specificity | Precision | Accuracy |
|---|---|---|---|---|---|---|---|---|---|
| DeNovoCNN | 63 | 52 | 11 | 98 | 14 | 78.79 | 89.91 | 82.54 | 85.71 |
| DeNovoGear-0.5 | NA | NA | NA | NA | NA | NA | NA | NA | NA |
| DeNovoGear-0.9 | NA | NA | NA | NA | NA | NA | NA | NA | NA |
| DeepTrio (DeepVariantWGS, BQSR) | 53 | 42 | 11 | 98 | 24 | 63.64 | 89.91 | 79.25 | 80.0 |
| DeepTrio (DeepVariantWGS, no BQSR) | 42 | 37 | 5 | 104 | 29 | 56.06 | 95.41 | 88.1 | 80.57 |
| DeepTrio (DeepVariant_unfiltered, BQSR) | 71 | 45 | 26 | 83 | 21 | 68.18 | 76.15 | 63.38 | 73.14 |
| DeepTrio (DeepVariant_unfiltered, no BQSR) | 65 | 47 | 18 | 91 | 19 | 71.21 | 83.49 | 72.31 | 78.86 |
| GATK | 84 | 54 | 30 | 79 | 12 | 81.82 | 72.48 | 64.29 | 76.0 |
| GATK_HC | 72 | 52 | 20 | 89 | 14 | 78.79 | 81.65 | 72.22 | 80.57 |
| In-house | 91 | 48 | 43 | 66 | 18 | 72.73 | 60.55 | 52.75 | 65.14 |

c) Deletions

| Tool | Calls | TP | FP | TN | FN | Sensitivity/ Recall | Specificity | Precision | Accuracy |
|---|---|---|---|---|---|---|---|---|---|
| DeNovoCNN | 112 | 100 | 12 | 87 | 47 | 68.03 | 87.88 | 89.29 | 76.02 |
| DeNovoGear-0.5 | 22 | 13 | 9 | 90 | 134 | 8.84 | 90.91 | 59.09 | 41.87 |
| DeNovoGear-0.9 | 20 | 13 | 7 | 92 | 134 | 8.84 | 92.93 | 65.0 | 42.68 |
| DeepTrio (DeepVariantWGS, BQSR) | 66 | 60 | 6 | 93 | 87 | 40.82 | 93.94 | 90.91 | 62.2 |
| DeepTrio (DeepVariantWGS, no BQSR) | 61 | 52 | 9 | 90 | 95 | 35.37 | 90.91 | 85.25 | 57.72 |
| DeepTrio | 78 | 68 | 10 | 89 | 79 | 46.26 | 89.9 | 87.18 | 63.82 |

| | | | | | | | | | |
|---|---|---|---|---|---|---|---|---|---|
| (DeepVariant_unfiltered, BQSR) | | | | | | | | | |
| DeepTrio (DeepVariant_unfiltered, no BQSR) | 77 | 69 | 8 | 91 | 78 | 46.94 | 91.92 | 89.61 | 65.04 |
| GATK | 142 | 109 | 33 | 66 | 38 | 74.15 | 66.67 | 76.76 | 71.14 |
| GATK_HC | 127 | 104 | 23 | 76 | 43 | 70.75 | 76.77 | 81.89 | 73.17 |
| In-house | 126 | 101 | 25 | 74 | 46 | 68.71 | 74.75 | 80.16 | 71.14 |

d) Total (substitutions + insertions + deletions) on chromosome 20

| **Tool** | **Calls** | **TP** | **FP** | **TN** | **FN** | **Sensitivity/ Recall** | **Specificity** | **Precision** | **Accuracy** |
|---|---|---|---|---|---|---|---|---|---|
| DeNovoCNN | 32 | 32 | 0 | 16 | 9 | 78.05 | 100.0 | 100.0 | 84.21 |
| DeNovoGear-0.5 | 40 | 34 | 6 | 10 | 7 | 82.93 | 62.5 | 85.0 | 77.19 |
| DeNovoGear-0.9 | 35 | 33 | 2 | 14 | 8 | 80.49 | 87.5 | 94.29 | 82.46 |
| DeepTrio (DeepVariantWGS, BQSR) | 3 | 3 | 0 | 16 | 38 | 7.32 | 100.0 | 100.0 | 33.33 |
| DeepTrio (DeepVariantWGS, no BQSR) | 4 | 4 | 0 | 16 | 37 | 9.76 | 100.0 | 100.0 | 35.09 |
| DeepTrio (DeepVariant_unfiltered, BQSR) | 35 | 33 | 2 | 14 | 8 | 80.49 | 87.5 | 94.29 | 82.46 |
| DeepTrio (DeepVariant_unfiltered, no BQSR) | 36 | 34 | 2 | 14 | 7 | 82.93 | 87.5 | 94.44 | 84.21 |
| GATK | 38 | 37 | 1 | 15 | 4 | 90.24 | 93.75 | 97.37 | 91.23 |
| GATK_HC | 37 | 36 | 1 | 15 | 5 | 87.8 | 93.75 | 97.3 | 89.47 |
| In-house | 41 | 35 | 6 | 10 | 6 | 85.37 | 62.5 | 85.37 | 78.95 |

**Supplementary Table 7. Comparison on manually curated GIAB dataset.** Every row shows different statistics and performance metrics for DeNovoCNN, DeNovoGear with the probability threshold of 0.5 and 0.9, DeepTrio with different settings, GATK and high quality GATK, and our in-house tool. The first column shows the name of the tool, the next five columns show the number of total calls, the number of true positive, false positive, true negative and false negative calls respectively based on the manually curated GIAB reference DNMs. The next four columns show sensitivity (recall), specificity, precision and accuracy. All statistics are shown for substitutions (a), insertions (b), deletions (c), and all (d) variants.

a) Substitutions

| Tool | Calls | TP | FP | TN | FN | Sensitivity/ Recall | Specificity | Precision | Accuracy |
|---|---|---|---|---|---|---|---|---|---|
| DeNovoCNN | 1058 | 976 | 82 | 463 | 19 | 98.09 | 84.95 | 92.25 | 93.44 |
| DeNovoGear-0.5 | 1324 | 984 | 340 | 205 | 11 | 98.89 | 37.61 | 74.32 | 77.21 |
| DeNovoGear-0.9 | 1141 | 972 | 169 | 376 | 23 | 97.69 | 68.99 | 85.19 | 87.53 |
| DeepTrio (DeepVariantWGS, BQSR) | 91 | 60 | 31 | 514 | 935 | 6.03 | 94.31 | 65.93 | 37.27 |
| DeepTrio (DeepVariantWGS, no BQSR) | 165 | 140 | 25 | 520 | 855 | 14.07 | 95.41 | 84.85 | 42.86 |
| DeepTrio (DeepVariant_unfiltered, BQSR) | 1058 | 949 | 109 | 436 | 46 | 95.38 | 80.0 | 89.7 | 89.94 |
| DeepTrio (DeepVariant_unfiltered, no BQSR) | 1080 | 951 | 129 | 416 | 44 | 95.58 | 76.33 | 88.06 | 88.77 |
| GATK | 1112 | 951 | 161 | 384 | 44 | 95.58 | 70.46 | 85.52 | 86.69 |
| GATK_HC | 1058 | 938 | 120 | 425 | 57 | 94.27 | 77.98 | 88.66 | 88.51 |
| In-house | 1076 | 969 | 107 | 438 | 26 | 97.39 | 80.37 | 90.06 | 91.36 |

b) Insertions

| Tool | Calls | TP | FP | TN | FN | Sensitivity/ Recall | Specificity | Precision | Accuracy |
|---|---|---|---|---|---|---|---|---|---|

| | | | | | | | | | |
|---|---|---|---|---|---|---|---|---|---|
| DeNovoCNN | 63 | 50 | 13 | 98 | 4 | 92.59 | 88.29 | 79.37 | 89.7 |
| DeNovoGear-0.5 | NA | NA | NA | NA | NA | NA | NA | NA | NA |
| DeNovoGear-0.9 | NA | NA | NA | NA | NA | NA | NA | NA | NA |
| DeepTrio (DeepVariantWGS, BQSR) | 53 | 41 | 12 | 99 | 13 | 75.93 | 89.19 | 77.36 | 84.85 |
| DeepTrio (DeepVariantWGS, no BQSR) | 42 | 37 | 5 | 106 | 17 | 68.52 | 95.5 | 88.1 | 86.67 |
| DeepTrio (DeepVariant_unfiltered, BQSR) | 71 | 44 | 27 | 84 | 10 | 81.48 | 75.68 | 61.97 | 77.58 |
| DeepTrio (DeepVariant_unfiltered, no BQSR) | 65 | 47 | 18 | 93 | 7 | 87.04 | 83.78 | 72.31 | 84.85 |
| GATK | 84 | 53 | 31 | 80 | 1 | 98.15 | 72.07 | 63.1 | 80.61 |
| GATK_HC | 72 | 51 | 21 | 90 | 3 | 94.44 | 81.08 | 70.83 | 85.45 |
| In-house | 91 | 47 | 44 | 67 | 7 | 87.04 | 60.36 | 51.65 | 69.09 |

c) Deletions

| Tool | Calls | TP | FP | TN | FN | Sensitivity/ Recall | Specificity | Precision | Accuracy |
|---|---|---|---|---|---|---|---|---|---|
| DeNovoCNN | 112 | 48 | 64 | 103 | 9 | 84.21 | 61.68 | 42.86 | 67.41 |
| DeNovoGear-0.5 | 22 | 7 | 15 | 152 | 50 | 12.28 | 91.02 | 31.82 | 70.98 |
| DeNovoGear-0.9 | 20 | 7 | 13 | 154 | 50 | 12.28 | 92.22 | 35.0 | 71.88 |
| DeepTrio (DeepVariantWGS, BQSR) | 66 | 27 | 39 | 128 | 30 | 47.37 | 76.65 | 40.91 | 69.2 |
| DeepTrio (DeepVariantWGS, no BQSR) | 61 | 22 | 39 | 128 | 35 | 38.6 | 76.65 | 36.07 | 66.96 |
| DeepTrio (DeepVariant_unfiltered, BQSR) | 78 | 30 | 48 | 119 | 27 | 52.63 | 71.26 | 38.46 | 66.52 |
| DeepTrio (DeepVariant_unfiltered, no | 77 | 31 | 46 | 121 | 26 | 54.39 | 72.46 | 40.26 | 67.86 |

| | | | | | | | | | |
|---|---|---|---|---|---|---|---|---|---|
| BQSR) | | | | | | | | | |
| GATK | 142 | 47 | 95 | 72 | 10 | 82.46 | 43.11 | 33.1 | 53.12 |
| GATK_HC | 127 | 45 | 82 | 85 | 12 | 78.95 | 50.9 | 35.43 | 58.04 |
| In-house | 126 | 46 | 80 | 87 | 11 | 80.7 | 52.1 | 36.51 | 59.38 |

d) Total (substitutions + insertions + deletions)

| **Tool** | **Calls** | **TP** | **FP** | **TN** | **FN** | **Sensitivity/ Recall** | **Specificity** | **Precision** | **Accuracy** |
|---|---|---|---|---|---|---|---|---|---|
| DeNovoCNN | 1233 | 1074 | 159 | 664 | 32 | 97.11 | 80.68 | 87.1 | 90.1 |
| DeNovoGear-0.5 | 1346 | 991 | 355 | 468 | 115 | 89.6 | 56.87 | 73.63 | 75.64 |
| DeNovoGear-0.9 | 1161 | 979 | 182 | 641 | 127 | 88.52 | 77.89 | 84.32 | 83.98 |
| DeepTrio (DeepVariantWGS, BQSR) | 210 | 128 | 82 | 741 | 978 | 11.57 | 90.04 | 60.95 | 45.05 |
| DeepTrio (DeepVariantWGS, no BQSR) | 268 | 199 | 69 | 754 | 907 | 17.99 | 91.62 | 74.25 | 49.4 |
| DeepTrio (DeepVariant_unfiltered, BQSR) | 1207 | 1023 | 184 | 639 | 83 | 92.5 | 77.64 | 84.76 | 86.16 |
| DeepTrio (DeepVariant_unfiltered, no BQSR) | 1222 | 1029 | 193 | 630 | 77 | 93.04 | 76.55 | 84.21 | 86.0 |
| GATK | 1338 | 1051 | 287 | 536 | 55 | 95.03 | 65.13 | 78.55 | 82.27 |
| GATK_HC | 1257 | 1034 | 223 | 600 | 72 | 93.49 | 72.9 | 82.26 | 84.71 |
| In-house | 1293 | 1062 | 231 | 592 | 44 | 96.02 | 71.93 | 82.13 | 85.74 |

**Supplementary Table 8. The results of the Sanger/IonTorrent validations on the 20 WES trios based on high quality calls of the tools.** Every row shows different statistics and performance metrics for DeNovoCNN, high quality GATK and GATK, DeNovoGear with a probability threshold of 0.9, our in-house tool, and DeepTrio with various different settings. The first column shows the name of the tool, the next five columns show the number of total calls, the number of true positive, false positive, true negative and false negative calls respectively based on the Sanger/IonTorrent validations. The next four columns show sensitivity (recall), specificity, precision and accuracy.

| Tool | Calls | TP | FP | TN | FN | Sensitivity/ Recall | Specificity | Precision | Accuracy |
|---|---|---|---|---|---|---|---|---|---|
| DeNovoCNN | 30 | 19 | 11 | 1843 | 0 | 100.0 | 99.41 | 63.33 | 99.41 |
| GATK_HC | 37 | 15 | 22 | 1832 | 4 | 78.95 | 98.81 | 40.54 | 98.61 |
| GATK | 38 | 15 | 23 | 1831 | 4 | 78.95 | 98.76 | 39.47 | 98.56 |
| DeNovoGear-0.9 | 84 | 15 | 69 | 1785 | 4 | 78.95 | 96.28 | 17.86 | 96.1 |
| In-house | 45 | 17 | 28 | 1826 | 2 | 89.47 | 98.49 | 37.78 | 98.4 |
| DeepTrio (DeepVariant_unfiltered, BQSR) | 1457 | 17 | 1440 | 414 | 2 | 89.47 | 22.33 | 1.17 | 23.01 |
| DeepTrio (DeepVariant_unfiltered, no BQSR) | 1490 | 16 | 1474 | 380 | 3 | 84.21 | 20.5 | 1.07 | 21.14 |
| DeepTrio (DeepVariantWES, BQSR) | 685 | 11 | 674 | 1180 | 8 | 57.89 | 63.65 | 1.61 | 63.59 |
| DeepTrio (DeepVariantWES, no BQSR) | 672 | 11 | 661 | 1193 | 8 | 57.89 | 64.35 | 1.64 | 64.28 |

**Supplementary Table 9. SolveRD dataset description.** The table shows the number of samples and average coverage for every sequencing kit (in rows) used in SolveRD dataset.

| **Sequencing kit** | **Number of samples** | **Average coverage** |
|---|---|---|
| Agilent_SureSelect_v3_51Mb | 8 | 79.51 |
| Illumina_TruSeq_v1.2_45Mb | 8 | 72.11 |
| Agilent_SureSelect_v7_36Mb | 12 | 112.25 |
| Agilent_SureSelect_v2_46Mb | 18 | 86.3 |
| Twist_Bioscience_Twist_Human_RefSeq_Panel | 21 | 67.8 |
| Agilent_SureSelect_CRE_V2_67Mb | 22 | 107.2 |
| Illumina_NexteraExpandedExome_62Mb | 23 | 112.8 |
| Nimblegen_SeqCapEZExome_v3_64Mb | 29 | 62.75 |
| Agilent_SureSelect_CRE_V1_54Mb | 33 | 100.78 |
| Agilent_SureSelect_v5_50Mb | 43 | 109.7 |
| Broad_Custom_v1_60Mb | 45 | 82.63 |
| Agilent_SureSelect_v6_60Mb | 51 | 122.13 |
| Nimblegen_SeqCapEZMedExomePlusMT_47Mb | 66 | 57.57 |
| Agilent_SureSelect_v4_51Mb | 78 | 75.26 |
| Illumina_NexteraExome_37Mb | 94 | 93.63 |
| **Total** | **551** | **89.49** |

**Supplementary Table 10. Validations on the 7 WGS trios using Pacbio LRS based on high quality calls.** Every row shows different statistics and performance metrics for DeNovoCNN, high quality GATK and GATK, and DeepTrio with two different settings. The first column shows the name of the tool, the next five columns show the number of total calls, the number of true positive, false positive, true negative and false negative calls respectively based on DNMs obtained from Pacbio HiFi LRS. The next four columns show sensitivity (recall), specificity, precision and accuracy.

| Tool | Calls | TP | FP | TN | FN | Sensitivity/ Recall | Specificity | Precision | Accuracy |
|---|---|---|---|---|---|---|---|---|---|
| DeNovoCNN | 1238 | 509 | 729 | 3458 | 26 | 95.14 | 82.59 | 41.11 | 84.01 |
| GATK_HC | 2567 | 471 | 2096 | 2091 | 64 | 88.04 | 49.94 | 18.35 | 54.26 |
| GATK | 3575 | 475 | 3100 | 1087 | 60 | 88.79 | 25.96 | 13.29 | 33.08 |
| DeepTrio (DeepVariant_unfiltered) | 1223 | 456 | 767 | 3420 | 79 | 85.23 | 81.68 | 37.29 | 82.08 |
| DeepTrio (DeepVariantWGS) | 413 | 97 | 316 | 3871 | 438 | 18.13 | 92.45 | 23.49 | 84.03 |

**Supplementary Table 11. DeNovoCNN calls on mosaic variants.** Every row shows the probability of DeNovoCNN for a mosaic variant found in a corresponding gene and location, supported with information about mosaic variants, such as number of reads, number of alternative reads, VAF, and whether the variant was confirmed using Sanger validations.

| DeNovoCNN probability | Gene | Chr change (Hg19) | reads | variation reads | VAF, % | Sanger validation |
|---|---|---|---|---|---|---|
| 0.784 | INA | Chr10(GRCh37):g.105046853T>C | 33 | 5 | 15.15 | - |
| 0.685 | SMC1A | ChrX(GRCh37):g.53440234C>T | 71 | 11 | 15.49 | True |
| 0.212 | EFTUD2 | Chr17(GRCh37):g.42945655G>A | 82 | 18 | 21.95 | True |
| 0.987 | GABBR2 | Chr9(GRCh37):g.101133817C>T | 41 | 13 | 31.71 | True |
| 0.993 | SATB2 | Chr2(GRCh37):g.200188570del | 143 | 32 | 22.38 | - |
| 0.98 | TBR1 | Chr2(GRCh37):g.162273213G>T | 226 | 38 | 16.81 | True |
| 0.978 | ASXL1 | Chr20(GRCh37):g.31023717C>T; | 105 | 25 | 23.81 | True |
| 0.947 | KANSL1 | Chr17(GRCh37):g.44249038_44249042del | 329 | 53 | 16.11 | True |
| 0.883 | EEF1A2 | Chr20(GRCh37):g.62127262C>T | 143 | 23 | 16.08 | True |
| 0.781 | KCNT1 | Chr9(GRCh37):g.138660693C>T | 85 | 14 | 16.47 | - |
