## Supplementary figures and images for "DeNovoCNN: A deep learning approach to *de novo* variant calling in next generation sequencing data"

### 20_WES_comarison_FP

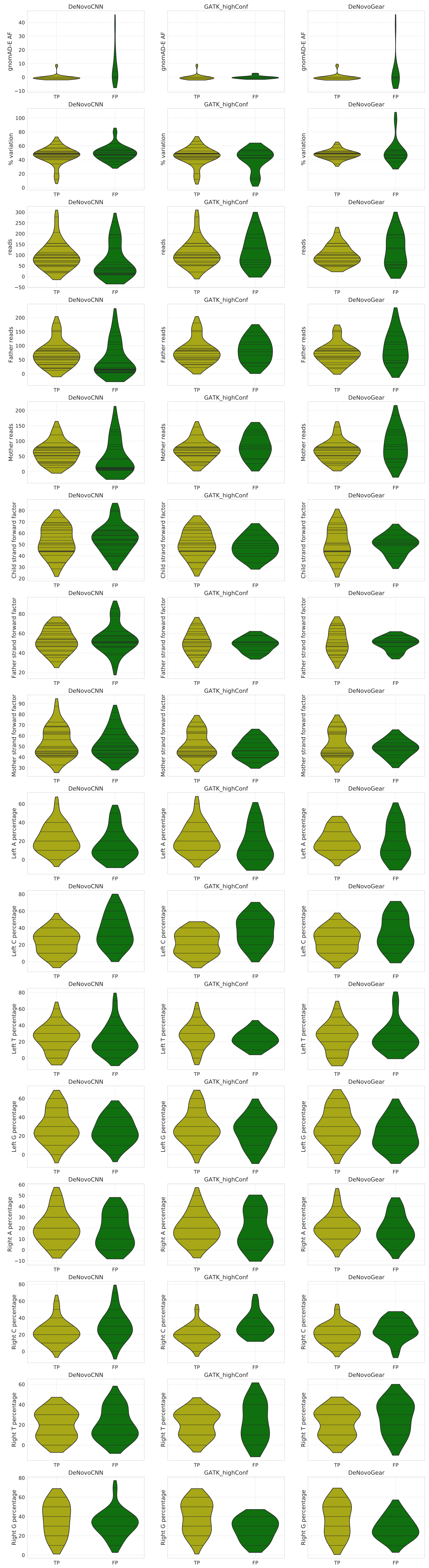
